# A meta-analysis of two high-risk prospective cohort studies reveals autism-specific transcriptional changes to chromatin, autoimmune, and environmental response genes in umbilical cord blood

**DOI:** 10.1101/486498

**Authors:** Charles E. Mordaunt, Bo Y. Park, Kelly M. Bakulski, Jason I. Feinberg, Lisa A. Croen, Christine Ladd-Acosta, Craig J. Newschaffer, Heather E. Volk, Sally Ozonoff, Irva Hertz-Picciotto, Janine M. LaSalle, Rebecca J. Schmidt, M. Daniele Fallin

## Abstract

**Background:** Autism spectrum disorder (ASD) is a neurodevelopmental disorder that affects more than 1% of children in the United States. ASD risk is thought to arise from both genetic and environmental factors, with the perinatal period as a critical window. Understanding early transcriptional changes in ASD would assist in clarifying disease pathogenesis and identifying biomarkers. However, little is known about umbilical cord blood gene expression profiles in babies later diagnosed with ASD compared to non-typically developing and non-ASD (Non-TD) or typically developing (TD) children.

**Methods:** Genome-wide transcript levels were measured by Affymetrix Human Gene 2.0 array in RNA from cord blood samples from both the Markers of Autism Risk in Babies--Learning Early Signs (MARBLES) and the Early Autism Risk Longitudinal Investigation (EARLI) high-risk pregnancy cohorts that enroll younger siblings of a child previously diagnosed with ASD. Younger siblings were diagnosed based on assessments at 36 months, and 59 ASD, 92 Non-TD, and 120 TD subjects were included. Using both differential expression analysis and weighted gene correlation network analysis, gene expression between ASD and TD, and between Non-TD and TD, was compared within each study and via meta-analysis.

**Results:** While cord blood gene expression differences comparing either ASD or Non-TD to TD did not reach genome-wide significance, 172 genes were nominally differentially-expressed between ASD and TD cord blood (log_2_(fold change) > 0.1, p < 0.01). These genes were significantly enriched for functions in xenobiotic metabolism, chromatin regulation, and systemic lupus erythematosus (FDR *q* < 0.05). In contrast, 66 genes were differentially-expressed between Non-TD and TD, including 8 genes that were also differentially-expressed in ASD. Gene coexpression modules were significantly correlated with demographic factors and cell type proportions.

**Limitations:** ASD-associated gene expression differences identified in this study are subtle, as cord blood is not the main affected tissue, it is composed of many cell types, and ASD is a heterogeneous disorder.

**Conclusions:** This is the first study to identify gene expression differences in cord blood specific to ASD. The results of this meta-analysis across two prospective pregnancy cohorts support involvement of environmental, immune, and epigenetic mechanisms in ASD etiology.

## Background

Autism spectrum disorder (ASD) is a neurodevelopmental disorder characterized by impaired social interaction and restricted and repetitive behaviors. Heritability of ASD risk has been well established with twin and family studies and is estimated at 52% [1–3]. While rare variants with large effects explain a relatively small proportion of all ASD cases, heritable common variants with individually minor effects contribute substantially to ASD risk [4]. Accumulating lines of evidence suggest that ASD arises from complex interactions between heterogeneous genetic and environmental risk factors. Gene expression levels are influenced by both genetic and environmental factors and determine the functional responses of cells and tissues. Postmortem brain gene expression studies have guided understanding of ASD pathophysiology and show evidence of changes in gene co-expression and enrichment in immune response and neuronal activity functions [5, 6]. Peripheral blood gene expression studies in children and adults using whole blood and in specific cell types (natural killer (NK) cell and lymphocytes) observed enrichment of immune and inflammatory processes in differential gene expression associated with ASD [7, 8]. Recent efforts have been focused on identifying how genetic risk factors converge into one or more unifying pathways and pathophysiological mechanisms [9, 10]. Yet, the majority of this work to date relies on post-mortem or post-symptom timing of sample collection, rather than prospective assessment of gene expression.

Converging evidence suggests that most of the changes in the brain associated with ASD are initiated during prenatal brain development [11, 12], but the complete nature of these changes remain unknown. Umbilical cord blood captures fetal blood as well as the exchanges across the feto-placental unit and provides a distinct insight into prenatal development. A unique cell mixture is represented in umbilical cord blood, including hematopoietic stem cells, B cells, NK cells, T cells, monocytes, granulocytes and nucleated red blood cells [13]. Cord blood gene expression would reflect the immune response as well as endocrine and cellular communication essential for fetal development near the time of birth.

While several studies have previously examined child blood gene expression differences in ASD [8, 14–20], this is the first study to take advantage of cord blood samples collected from two prospective studies (Markers of Autism Risk in Babies--Learning Early Signs (MARBLES) and the Early Autism Risk Longitudinal Investigation (EARLI)) in order to assess the perinatal transcriptional changes that precede ASD diagnosis in high-risk children [21, 22]. The subjects in this study are all siblings of children with ASD and thus have a 13-fold increased risk for ASD compared to the general population [23]. They are also at a higher risk for non-typical neurodevelopment, including deficits in attention and behavior. We measured cord blood gene expression levels using the Affymetrix Human Gene 2.0 array and compared gene-level differential expression, gene set enrichment, and gene coexpression networks across ASD, non-typically developing (Non-TD) and neurotypical children (Additional file 1: Fig. S1). Study-level results were then combined in a meta-analysis to investigate cord blood transcriptional dysregulation in ASD.

## Methods

### Sample population and biosample collection

#### MARBLES

The MARBLES study recruits Northern California mothers from lists of children receiving services through the California Department of Developmental Services who have a child with confirmed ASD and are planning a pregnancy or are pregnant with another child. Inclusion criteria for the study were: 1) mother or father has one or more biological child(ren) with ASD; 2) mother is 18 years or older; 3) mother is pregnant; 4) mother speaks, reads, and understands English sufficiently to complete the protocol and the younger sibling will be taught to speak English; 5) mother lives within 2.5 hours of the Davis/Sacramento region at time of enrollment. As described in more detail elsewhere [21], demographic, diet, lifestyle, environmental, and medical information were prospectively collected through telephone-assisted interviews and mailed questionnaires throughout pregnancy and the postnatal period. Mothers were provided with sampling kits for cord blood collection prior to delivery. MARBLES research staff made arrangements with obstetricians/midwives and birth hospital labor and delivery staff to assure proper sample collection and temporary storage. Infants received standardized neurodevelopmental assessments beginning at 6 months, as described below, and concluding at 3 years of age. For this study, all children actively enrolled by March 1, 2017 (*n* = 347) with umbilical cord blood collected in a PAXgene Blood RNA tube (*n* = 262, 76%) were included.

#### EARLI

The EARLI study is a high-risk pregnancy cohort that recruited and followed pregnant mothers who had an older child diagnosed with ASD through pregnancy, birth, and the first three years of life. EARLI families were recruited at four EARLI Network sites (Drexel/Children’s Hospital of Philadelphia; Johns Hopkins/Kennedy Krieger Institute; University of California (UC) Davis; and Kaiser Permanente Northern California) in three distinct US regions (Southeast Pennsylvania, Northeast Maryland, and Northern California). In addition to having a biological child with ASD confirmed by EARLI study clinicians, to be eligible mothers also had to communicate in English or Spanish and, at recruitment, meet the following criteria: be 18 years or older; live within two hours of a study site; and be < 29 weeks pregnant. The design of the EARLI study is described in more detail in Newschaffer et al. [22]. EARLI research staff made arrangements with obstetricians/midwives and birth hospital labor and delivery staff to ensure proper cord blood sample collection and temporary storage. The development of children born into the cohort was closely followed through age three years. For this study, 212 infants born into EARLI as a singleton birth and followed to one year of age were considered for inclusion. Of the 212 infants, 97 were excluded because they were either missing umbilical cord blood samples or outcome measures at 36 months, leaving a final sample of 115.

##### Diagnostic outcomes

In both studies, development was assessed by trained, reliable examiners. Diagnostic assessments at three years included the gold standard Autism Diagnostic Observation Schedule (ADOS) [24, 25], the Autism Diagnostic Interview-Revised (ADI-R) [26] conducted with parents, and the Mullen Scales of Early Learning (MSEL) [27], a test of cognitive, language, and motor development. Participants were classified into one of three outcome groups, ASD, typically developing (TD), and Non-TD, based on a previously published algorithm that uses ADOS and MSEL scores [28, 29]. Children with ASD outcomes had scores over the ADOS cutoff and met DSM-5 criteria for ASD. The Non-TD group was defined as children with low MSEL scores (i.e., two or more MSEL subscales that are more than 1.5 standard deviations (SD) below average or at least one MSEL subscale that was more than 2 SD below average), elevated ADOS scores (i.e., within 3 points of the ASD cutoff), or both. Children with TD outcomes had all MSEL scores within 2 SD and no more than one MSEL subscale 1.5 SD below the normative mean and scores on the ADOS at least three or more points below the ASD cutoff.

##### Demographic characteristics

In both the MARBLES and EARLI studies, demographic information was prospectively collected through in-person and telephone-assisted interviews and mailed questionnaires throughout pregnancy and the postnatal period. Cotinine was measured in maternal urine during pregnancy, and maternal smoking was identified if the concentration of cotinine was > 50 ng/mL [30]. Within each study, demographic characteristics were stratified by diagnostic outcome and compared using Fisher’s exact test for categorical variables and one-way ANOVA for continuous variables.

##### RNA isolation and expression assessment

In both MARBLES and EARLI, umbilical cord blood was collected at the time of birth in PAXgene Blood RNA Tubes with the RNA stabilization reagent (BD Biosciences) and stored at −80°C. RNA isolation was performed with the PAXgene Blood RNA Kit (Qiagen) following the manufacturer’s protocol. RNA from 236 (90%) of the 262 MARBLES PAXgene blood samples and all of the EARLI PAXgene blood samples met quality control standards (RIN ≥ 7.0 and concentration ≥ 35ng/uL) and volume requirements. Total RNA was converted to cDNA and *in vitro* transcribed to biotin-labeled cRNA, which was hybridized to Human Gene 2.0 Affymetrix microarray chips by the Johns Hopkins Sequencing and Microarray core. EARLI and MARBLES samples were measured separately and in multiple batches within each study. The manufacturer’s protocol was followed for all washing, staining and scanning procedures. The raw fluorescence data (in Affymetrix CEL file format) with one perfect match and one mismatched probe in each set were analyzed using oligo package in R.

##### Data preprocessing

Within each study, signal distribution was first assessed in perfect-match probe intensity and Robust Multi-Chip Average (RMA) normalized data [31]. During the quality control step, we identified outliers using the arrayQualityMetrics and oligo R packages [32, 33]. Outliers were excluded based on loading in principal component 1, the Kolmogorov-Smirnov test, median normalized unscaled standard error, and the sum of the distances to all other arrays. For the MARBLES study, 3 outlier samples were identified and excluded, and another 71 children had not yet received a diagnosis by April 12, 2018 so were excluded; 162 samples were normalized using RMA. For the EARLI study, 6 outliers were identified and excluded, then 109 samples were normalized using RMA. Probes were annotated at the transcript level using the pd.hugene.2.0.st R package [34], and those assigned to a gene (36,459 probes) were used in subsequent analyses.

##### Surrogate variable analysis

Surrogate variable analysis (SVA) was used to estimate and adjust for unmeasured environmental, demographic, cell type proportion, and technical factors that may have substantial effects on gene expression using the SVA R package [35, 36]. 21 surrogate variables were detected in normalized expression data from MARBLES subjects for both the ASD versus TD and Non-TD versus TD comparisons. Specific factors associated with surrogate variables in MARBLES using linear regression included array batch, sex, maternal BMI, gestational age, delivery method, child ethnicity, and maternal education (False Discovery Rate (FDR) *q* < 0.1, Additional file 1: Fig. S2a). In normalized expression data from EARLI subjects, 11 surrogate variables were detected for the ASD versus TD comparison, which were associated with sex, birthweight, gestational age, and paternal age (FDR *q* < 0.1, Additional file 1: Fig. S3a). 12 surrogate variables were detected for the Non-TD versus TD comparison, which were associated with sex and gestational age (FDR *q* < 0.1, Additional file 1: Fig. S4a).

Proportion of variance in expression of each gene explained by each surrogate variable was determined using the variancePartition R package [37]. Median variance explained by each surrogate variable ranged from 0.3% to 5.6% in MARBLES (Additional file 1: Fig. S2b). In EARLI, median variance explained by each surrogate variable ranged from 0.8% to 7.1% for the ASD versus TD comparison and 0.5% to 7.2% for the Non-TD versus TD comparison (Additional file 1: Fig. S3b, S4b).

##### Differential gene expression

Differential expression was determined using the limma package in R with diagnosis and all surrogate variables included in the linear model [38] (Additional file 1: Fig. S5, S6). ASD versus TD and Non-TD versus TD differential expression results were extracted from a single model with three levels for diagnosis for MARBLES, while two pairwise models were used for EARLI, although this did not affect the results (1 vs 2 model meta-analysis fold change ASD vs TD Pearson’s r = 0.97, Non-TD vs TD Pearson’s r = 0.99). Fold change and standard error from each study were input into the METAL command-line tool for meta-analysis using the standard error analysis scheme with genomic control correction [39]. In this approach, the fold changes from each study are weighted using the inverse of the standard error. Using the meta-analyzed data, differential probes were then identified as those with a nominal *p*-value < 0.01 and an average absolute log_2_(fold change) > 0.1.

##### Gene overlap analysis

Gene overlap analysis by Fisher’s exact test was performed using the GeneOverlap R package [40]. Gene symbols annotated to differentially-expressed probes were compared to autism-related or blood cell-type associated gene lists [41] for overlap relative to all genes annotated to probes on the array. Genes with variation previously associated with autism were obtained from the Simons Foundation Autism Research Initiative (SFARI) Gene database and a recent genome-wide association study meta-analysis [42, 43], while genes with expression previously associated with autism were obtained from multiple previous reports [6, 8, 44, 45]. Significant overlaps were those with an FDR *q*-value < 0.05.

##### Overrepresentation enrichment analysis

Differential probes identified during meta-analysis were converted to Entrez gene IDs using the biomaRt R package [46]. Functional enrichment of only differential probes by hypergeometric test was relative to all probes on the array and was performed using the WebGestalt online tool with default parameters for the overrepresentation enrichment analysis method [47]. Enrichment databases included WebGestalt defaults and also a custom database of recently evolved genes obtained from [48]. WebGestalt default databases queried included Gene Ontology, KEGG, WikiPathways, Reactome, PANTHER, MSigDB, Human Phenotype Ontology, DisGeNET, OMIM, PharmGKB, and DrugBank. Significant enrichments were those with an FDR *q*-value < 0.05.

##### Gene set enrichment analysis (GSEA)

All probes included in the analysis were ranked using meta-analysis log_2_(fold change) and input into the WebGestalt online tool using default parameters for the GSEA method [47]. GSEA assesses whether genes in biologically-predefined sets occur toward the top or bottom of a ranked list of all examined genes more than expected by chance [49]. GSEA calculates an enrichment score normalized to the set size to estimate the extent of non-random distribution of the predefined gene set, and it then tests the significance of the enrichment with a permutation test. Enrichment databases included WebGestalt defaults (see above). Significant gene sets were called as those with an FDR *q*-value < 0.05.

##### Weighted gene correlation network analysis (WGCNA)

WGCNA was performed using the WGCNA R package [50]. RMA-normalized expression data were adjusted for batch using ComBat due to large batch effects in MARBLES [36]. Samples were clustered with hierarchical clustering using the average method and excluded using static tree cutting with cut height set to 100, resulting in one outlier being removed for each study [50]. Expression data were renormalized with RMA and adjusted for batch after removing outliers. Final samples for WGCNA included 59 ASD (41 MARBLES/19 EARLI), 91 Non-TD (44 MARBLES/47 EARLI), and 119 TD (76 MARBLES/43 EARLI). Signed topological overlap matrices (TOMs) were obtained separately for each study in a single block using the biweight midcorrelation with the maximum percentile for outliers set to 0.1 and the soft-thresholding power set to 10. Study-specific TOMs were calibrated using full quantile normalization, and a consensus TOM was calculated as the parallel minimum of the study-specific TOMs. Modules were identified using dynamic hybrid tree cutting with deepSplit set to 4, and modules with a dissimilarity < 0.1 were merged. Module hub probes were determined as the probe in each module with the highest module membership. Study-specific module eigengenes were correlated with demographic factors or estimated cell type proportions using the biweight midcorrelation with the maximum percentile for outliers set to 0.1 and including only pairwise-complete observations. Study-specific correlation Z-scores were combined in a meta-analysis using Stouffer’s method with weights given by the square root of the sample *n* [51]. P-values were adjusted for all comparisons using the FDR method. Significant correlations were called as those with an FDR *q*-value < 0.05.

##### Cell type proportion deconvolution

Estimation of cell type proportions was performed using CIBERSORT [41]. Final samples for cell type deconvolution were the same as for WGCNA: 59 ASD (41 MARBLES/19 EARLI), 91 Non-TD (44 MARBLES/47 EARLI), and 119 TD (76 MARBLES/43 EARLI). RMA-normalized expression data was adjusted for batch due to large batch effects in MARBLES. To correspond with identifiers used by CIBERSORT, array probes were matched to HUGO Gene Nomenclature Committee (HGNC) gene symbols using the biomaRt R package [46]. RMA-normalized expression data and the default LM22 adult blood signature genes file were input into the CIBERSORT web tool [41]. A similar signature genes file was not available for umbilical cord blood. Relative and absolute modes were run together, with 100 permutations and without quantile normalization. Deconvolution goodness of fit *p*-value was < 0.05 for all subjects.

Estimated cell type proportions were correlated with demographic factors within each study using the biweight midcorrelation with the maximum percentile for outliers set to 0.1 and including only pairwise-complete observations. Study-specific correlation Z-scores were combined in a meta-analysis using Stouffer’s method with weights given by the square root of the sample *n* [51]. P-values were adjusted for all comparisons using the FDR method. Significant correlations were called as those with an FDR *q*-value < 0.05.

## Results

### Study sample characteristics

MARBLES subjects in the final analysis included 41 ASD (30 male, 11 female), 44 Non-TD (27 male, 17 female), and 77 TD subjects (40 male, 37 female). Paternal age and gestational age were nominally associated with diagnostic group in MARBLES, with slightly increased paternal age and gestational age for the ASD subjects (paternal age *p* = 0.02, gestational age *p* = 0.04, Table 1). Other demographic characteristics were not associated with diagnostic group among MARBLES subjects. EARLI subjects in the final analysis included 18 ASD (13 male, 5 female), 48 Non-TD (23 male, 25 female), and 43 TD subjects (19 male, 24 female). Child race and ethnicity and home ownership were nominally associated with diagnostic group in EARLI (race and ethnicity *p* = 0.02, home ownership *p* = 0.01, Table 2). Specifically, the ASD group included a lower proportion of white subjects and a lower rate of home ownership. Other demographic characteristics were not associated with diagnostic group among EARLI subjects. In the meta-analysis, which combined both the MARBLES and EARLI studies, gene expression was analyzed in 271 subjects, including 120 TD, 59 ASD, and 92 Non-TD subjects.

**Table 1.** Demographic characteristics of children and their parents in the MARBLES study, stratified by child diagnosis.

|  | Child 36 Month Outcome |  |  | *p*-value<sup>a</sup> |
| --- | --- | --- | --- | --- |
|  | ASD<br>(*n* = 41) | Non-TD<br>(*n* = 44) | TD<br>(*n* = 77) |  |
| Child Male Gender, *n* (%) | 30 (73.2) | 27 (61.4) | 40 (51.9) | 0.08 |
| Child race and ethnicity, *n* (%) |  |  |  | 0.90 |
| White (European or Middle Eastern) | 19 (46.3) | 18 (40.9) | 38 (49.4) |  |
| Black/African-American | 2 (4.9) | 2 (4.5) | 1 (1.3) |  |
| Asian or Pacific Islander | 4 (9.8) | 3 (6.8) | 7 (9.1) |  |
| Hispanic White | 8 (19.5) | 11 (25) | 19 (24.7) |  |
| Hispanic Non-White | 4 (9.8) | 3 (6.8) | 6 (7.8) |  |
| Multiracial | 4 (9.8) | 7 (15.9) | 6 (7.8) |  |
| Mother age (years), *mean* (*SD*)<sup>b</sup> | 35.4 (5.1) | 34.1 (4.3) | 33.3 (4.9) | 0.09 |
| Father age (years), *mean* (*SD*)<sup>c</sup> | 38.7 (5.3) | 36.1 (4.8) | 35.7 (6.2) | **0.02** |
| Mother pre-pregnancy BMI, *mean* (*SD*)<sup>d</sup> | 27.9 (7.4) | 29.3 (8.3) | 26.1 (6.2) | 0.05 |
| Mother Type 1 Diabetes, *n* (%)<sup>e</sup> | 0 (0) | 0 (0) | 1 (1.3) | 1.00 |
| Mother Type 2 Diabetes, *n* (%)<sup>e</sup> | 0 (0) | 0 (0) | 1 (1.3) | 1.00 |
| Mother Gestational Diabetes, *n* (%)<sup>f</sup> | 12 (30) | 6 (14.3) | 11 (14.7) | 0.12 |
| Mother Hypertension, *n* (%)<sup>e</sup> | 1 (2.6) | 1 (2.3) | 1 (1.3) | 1.00 |
| Mother Preeclampsia, *n* (%)<sup>g</sup> | 2 (5.1) | 4 (10) | 2 (2.7) | 0.29 |
| Mother Smoking by Urine Cotinine, *n* (%)<sup>h</sup> | 4 (17.4) | 1 (4.3) | 1 (2.9) | 0.12 |
| Mother bachelor's degree +, *n* (%) | 18 (43.9) | 19 (43.2) | 40 (51.9) | 0.57 |
| Father bachelor's degree +, *n* (%) | 21 (51.2) | 19 (43.2) | 39 (50.6) | 0.69 |
| Own home, *n* (%)<sup>i</sup> | 21 (55.3) | 25 (58.1) | 48 (63.2) | 0.69 |
| Cesarean Delivery, *n* (%)<sup>j</sup> | 11 (26.8) | 17 (38.6) | 31 (41.3) | 0.28 |
| Gestational age (weeks), *mean* (*SD*) | 39.4 (1.1) | 39 (1.3) | 38.7 (1.4) | **0.04** |
| Birth weight (kg), *mean* (*SD*) | 3.5 (0.4) | 3.5 (0.4) | 3.4 (0.5) | 0.52 |

Autism spectrum disorder (ASD), Non-typically developing (Non-TD), Typically developing (TD);
<sup>a</sup> *p*-values from Fisher's exact test for categorical variables and one-way ANOVA for continuous variables; <sup>b</sup> Frequency missing = 1 ASD; <sup>c</sup> Frequency missing = 2 ASD, 1 TD; <sup>d</sup> Frequency missing = 1 ASD, 1 TD; <sup>e</sup> Frequency missing = 2 ASD, 1 Non-TD, 1 TD; <sup>f</sup> Frequency missing = 1 ASD, 2 Non-TD, 2 TD; <sup>g</sup> Frequency missing = 2 ASD, 4 Non-TD, 2 TD; <sup>h</sup> Frequency missing = 18 ASD, 21 Non-TD, 42 TD; <sup>i</sup> Frequency missing = 3 ASD, 1 Non-TD, 1 TD; <sup>j</sup> Frequency missing = 2 TD;

**Table 2.** Demographic characteristics of children and their parents in the EARLI study, stratified by child diagnosis.

|  | Child 36 Month Outcome |  |  | *p*-value<sup>a</sup> |
| --- | --- | --- | --- | --- |
|  | ASD<br>(*n* = 18) | Non-TD<br>(*n* = 48) | TD<br>(*n* = 43) |  |
| Child Male Gender, *n* (%) | 13 (72.2) | 23 (47.9) | 19 (44.2) | 0.13 |
| Child race and ethnicity, *n* (%)<sup>b</sup> |  |  |  | **0.02** |
| White (European or Middle Eastern) | 6 (35.3) | 23 (48.9) | 24 (55.8) |  |
| Black/African-American | 3 (17.6) | 8 (17) | 1 (2.3) |  |
| Asian or Pacific Islander | 2 (11.8) | 5 (10.6) | 0 (0) |  |
| Hispanic White | 2 (11.8) | 3 (6.4) | 3 (7) |  |
| Hispanic Non-White | 2 (11.8) | 7 (14.9) | 8 (18.6) |  |
| Multiracial | 2 (11.8) | 1 (2.1) | 7 (16.3) |  |
| Mother age (years), *mean* (*SD*) | 34.4 (4) | 33.8 (4.3) | 34.7 (5) | 0.60 |
| Father age (years), *mean* (*SD*) | 35.4 (6.8) | 35.7 (5) | 36.2 (6.4) | 0.88 |
| Mother pre-pregnancy BMI, *mean* (*SD*)<sup>c</sup> | 29.4 (7.1) | 27.4 (7) | 27.9 (7.2) | 0.60 |
| Mother Smoking by Urine Cotinine, *n* (%)<sup>d</sup> | 1 (6.3) | 1 (2.3) | 0 (0) | 0.30 |
| Mother bachelor's degree +, *n* (%) | 8 (44.4) | 28 (58.3) | 29 (67.4) | 0.26 |
| Father bachelor's degree +, *n* (%)<sup>e</sup> | 7 (38.9) | 28 (62.2) | 25 (58.1) | 0.24 |
| Own home, *n* (%) | 8 (44.4) | 31 (64.6) | 36 (83.7) | **0.01** |
| Cesarean Delivery, *n* (%)<sup>f</sup> | 9 (56.3) | 17 (39.5) | 12 (35.3) | 0.38 |
| Gestational age (weeks), *mean* (*SD*) | 39.2 (1.4) | 39.4 (1.5) | 39.1 (1.5) | 0.70 |
| Birth weight (kg), *mean* (*SD*)<sup>g</sup> | 3.5 (0.6) | 3.5 (0.6) | 3.5 (0.6) | 1.00 |

Autism spectrum disorder (ASD), Non-typically developing (Non-TD), Typically developing (TD);
<sup>a</sup> *p*-values from Fisher's exact test for categorical variables and one-way ANOVA for continuous variables; <sup>b</sup> Frequency missing = 1 ASD, 1 Non-TD; <sup>c</sup> Frequency missing = 1 Non-TD; <sup>d</sup> Frequency missing = 2 ASD, 4 Non-TD, 3 TD; <sup>e</sup> Frequency missing = 3 Non-TD; <sup>f</sup> Frequency missing = 2 ASD, 5 Non-TD, 9 TD; <sup>g</sup> Frequency missing = 1 TD

### ASD-associated differential gene expression in cord blood

We examined differential expression of single genes in cord blood samples in association with ASD diagnosis status at 36 months. In meta-analysis, no transcripts were differentially expressed at a conservative FDR q-value < 0.05. Under the thresholds of log_2_(fold change) > 0.1 and nominal *p*-value < 0.01, 172 transcripts were differentially expressed between ASD and TD cord blood (ASD *n* = 59, TD *n* = 120, Fig. 1a, Additional file 2: Table S1). Among these differential transcripts, 87 were upregulated and 85 were downregulated, and the median absolute log_2_(fold change) was 0.12. The differential transcript with the greatest absolute fold change was TUBB2A (log_2_(fold change) = 0.35, standard error = 0.12, p = 4.8E-3, Fig. 1b, Table 3). Additionally, the estimated fold changes for differentially-expressed genes were strongly correlated between the two studies (Pearson’s *r* = 0.80, *p* < 2.2E-16), although the fold changes of all transcripts were weakly correlated (Pearson’s r = 0.02, *p* = 4.6E-4, Additional File 1: Fig. S7a). Many of the differentially-expressed genes were noncoding or uncharacterized transcripts; however, the median expression of differentially-expressed genes was not lower than non-differentially-expressed genes on the array (MARBLES: differential = 4.70, non-differential =, *p* = 0.74; EARLI: differential = 4.34, non-differential = 4.19, *p* = 0.52; Additional file 1: Fig. S8).

**Figure 1.**
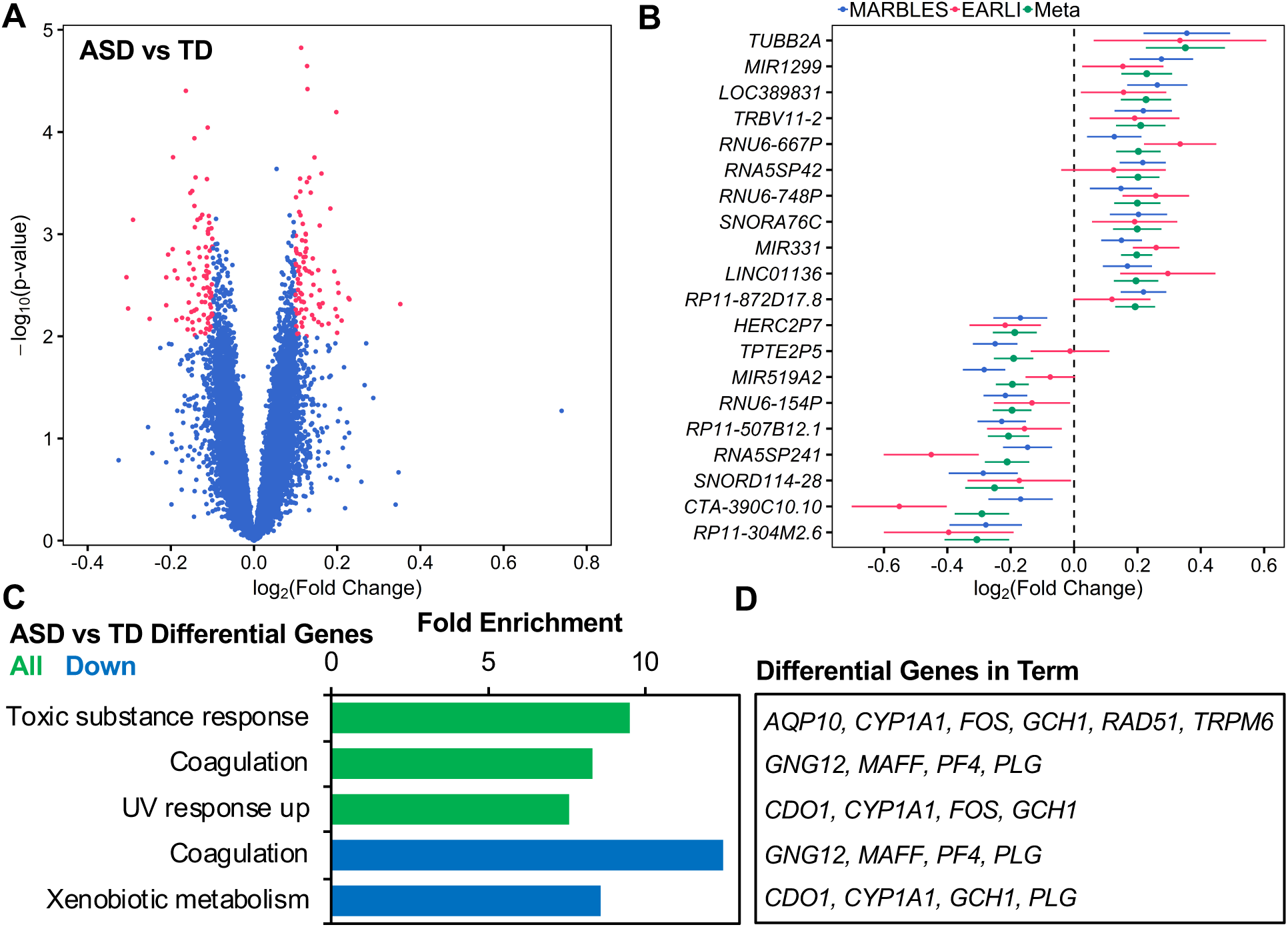
Identification and function of ASD-associated differentially-expressed genes in cord blood from two high-risk prospective studies. Gene expression in umbilical cord blood samples from subjects with typical development (*n* = 120, 59 male/61 female) or those diagnosed with ASD at age 36 months (*n* = 59, 43 male/16 female) was assessed by expression microarray. SVA was performed to control for technical and biological variables including sex and array batch. Differential expression analysis was carried out separately by study and combined in a meta-analysis. (A) Identification of 172 differentially-expressed genes in meta-analysis (178 probes, log_2_(fold change) > 0.1, p < 0.01). (B) Fold change in gene expression for top 20 differentially-expressed genes sorted by meta-analysis log_2_(fold change) and plotted for individual studies and meta-analysis. (C) ASD-associated differentially-expressed genes were analyzed for functional enrichment with WebGestalt using the hypergeometric test and compared to all genes annotated to the array. Significantly-enriched ontology terms are shown (FDR *q* < 0.05). (D) ASD-associated differentially-expressed genes belonging to significantly-enriched ontology terms are listed.

**Table 3.**
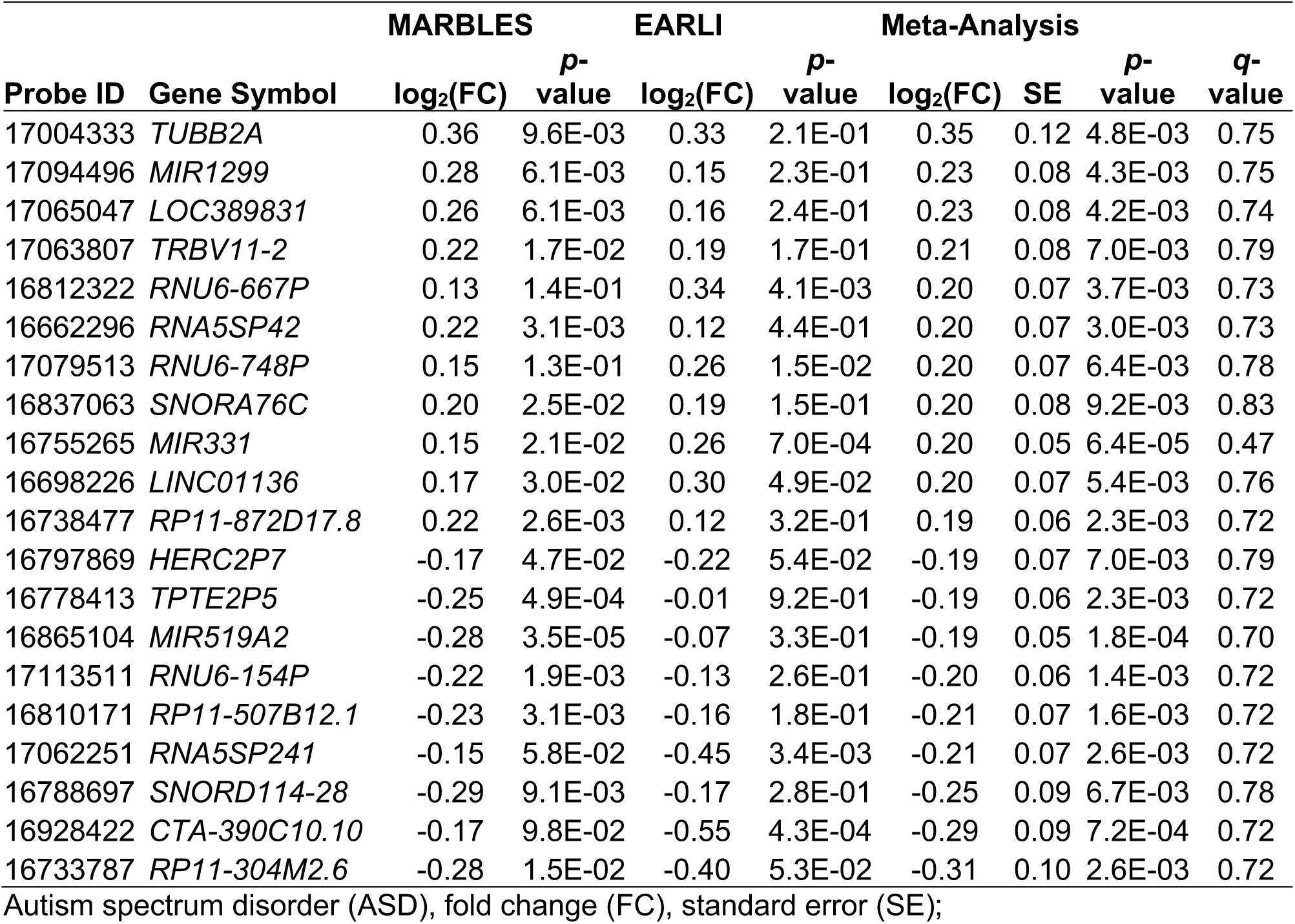
Top 20 ASD-associated differentially-expressed genes by log_2_(fold change) in meta-analysis.

Several differentially-expressed genes in cord blood overlapped with genes previously associated with ASD in genetic or gene expression studies, although the overlap was not statistically significant (FDR *q* > 0.05, Additional file 1: Fig. S9, Additional file 2: Table S2). Specifically, *SLC7A3*, *VSIG4*, and *MIR1226* have been associated with ASD in genetic studies [42, 52], while *SNORD104*, *OR2AG2*, and *DHX30* were differentially expressed in the same direction in ASD in gene expression studies [8, 44]. Further, *GFI1*, *GPR171*, *KIR2DL4, PTGIR*, and *TRPM6* were differentially expressed in ASD cord blood and are also differentially expressed in specific blood cell types, including natural killer cells and T cells, although a significant enrichment was not observed (*q* > 0.05, Additional file 1: Fig. S10, Additional file 2: Table S3) [41].

Overrepresentation enrichment analysis, which looks for overlap of only differentially-expressed genes with biologically-predefined gene lists, revealed that ASD differential transcripts were significantly enriched for functions in the response to toxic substances (*TRPM6*, *CYP1A1*, *FOS*, *GCH1*, *AC012476.1*, *RAD51*, and *AQP10*, fold enrichment = 9.5, *q* = 0.027) and ultraviolet radiation (*CDO1*, *CYP1A1*, *FOS*, and *GCH1*, fold enrichment = 7.6, *q* = 0.037, Fig. 1c, Additional file 2: Table S4). Both of these functional enrichments included the genes *CYP1A1, FOS*, and *GCH1*. Additionally, downregulated transcripts were enriched for functioning in blood coagulation (*GNG12*, *MAFF, PF4*, and *PLG,* fold enrichment = 12.5, *q* = 0.009) and xenobiotic metabolism (*CDO1*, *CYP1A1*, *GCH1*, and *PLG,* fold enrichment = 8.6, *q* = 0.019), but no significant enrichments were observed for upregulated transcripts alone.

Using fold changes to rank all transcripts for gene set enrichment analysis (GSEA), we observed significant enrichment for upregulation of gene sets involved in chromatin regulation (*q* < 0.05, Fig. 2, Additional file 2: Table S5). In other words, genes associated with chromatin regulation tended to be ranked toward the top of the distribution of fold change in ASD cord blood. Chromatin gene sets upregulated in ASD included DNA methylation (23 leading edge (LE) transcripts, normalized enrichment score (NES) = 2.16, *q* = 0.009), condensation of prophase chromosomes (24 LE transcripts, NES = 2.11, *q* = 0.007), nucleosome assembly (24 LE transcripts, NES = 1.96, *q* = 0.026), histone deacetylase (HDAC)-mediated deacetylation (30 LE transcripts, NES = 1.90, *q* = 0.040), and polycomb-repressive complex 2 (PRC2)-mediated methylation (22 LE transcripts, NES = 1.89, *q* = 0.002). Additionally, the gene set for the autoimmune disease systemic lupus erythematosus was significantly upregulated (45 LE transcripts, NES = 2.13, *q* = 0.003). Most of the genes associated with these sets compose a cluster of histone genes located at the 6p22.2 locus, which was also enriched (27 LE transcripts, NES = 2.15, *q* = 0.007). The above findings of differential expression across two prospective cohorts suggest transcriptional dysregulation in environmentally-responsive genes is present at birth in cord blood of high-risk subjects later diagnosed with ASD.

**Figure 2.**
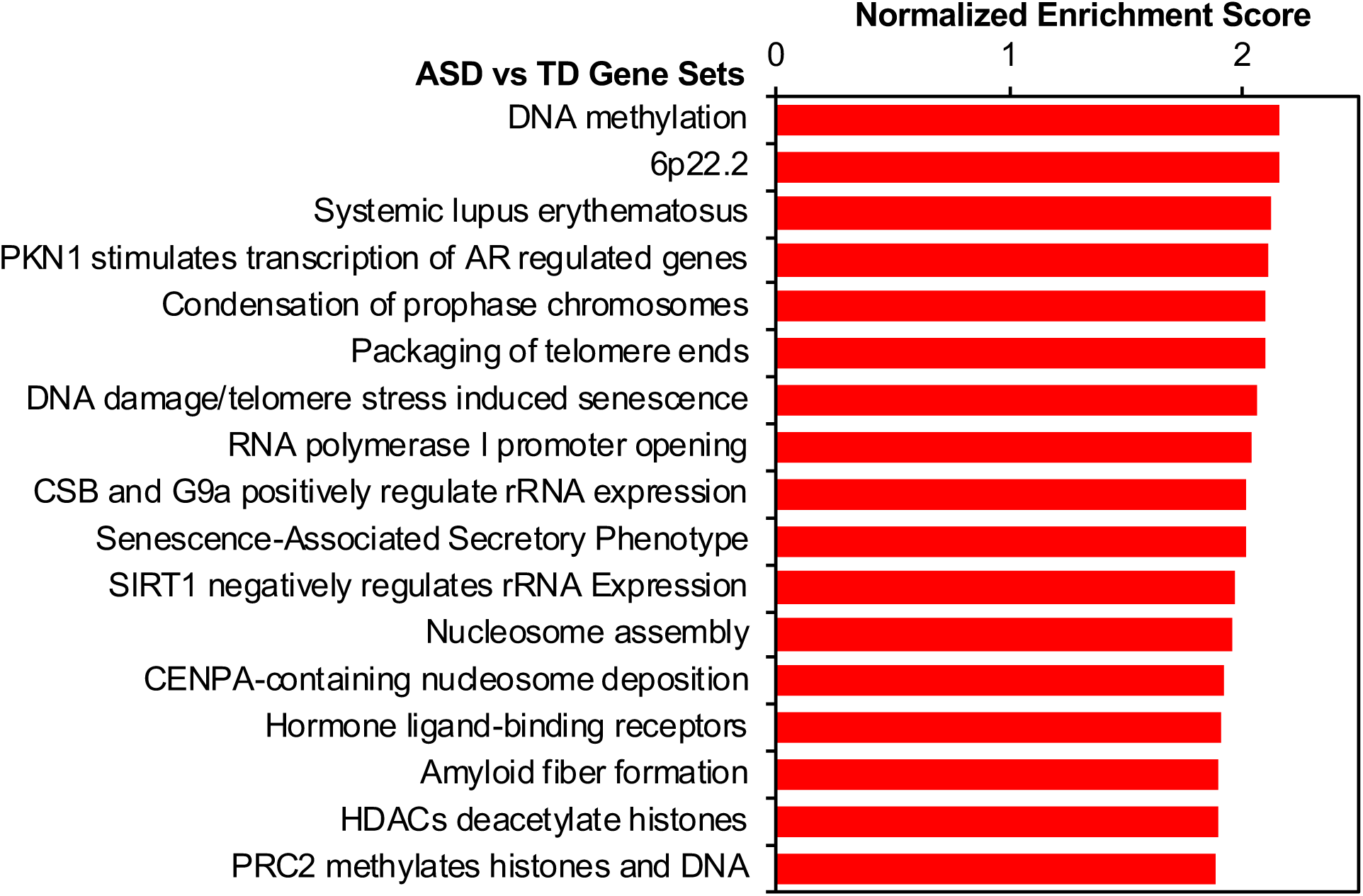
Chromatin and autoimmune gene sets are upregulated in cord blood from ASD subjects. Meta-analysis log_2_(fold change) for ASD versus TD gene expression was used to rank probes for gene set enrichment analysis (GSEA) with WebGestalt. GSEA assesses whether genes in biologically-defined sets occur toward the top or bottom of a ranked list more than expected by chance. Significantly-enriched gene sets are shown (FDR *q* < 0.05).

### Non-TD-associated differential gene expression in cord blood

To assess the specificity of ASD-associated transcriptional differences in cord blood, we also examined differential expression between cord blood samples from infants later classified as Non-TD compared to TD at 36 months. Meta-analysis results showed no transcripts differentially expressed at a conservative FDR q-value < 0.05. Under the thresholds of log_2_(fold change) > 0.1 and nominal *p*-value < 0.01, 66 transcripts were differential, with 38 upregulated and 28 downregulated (Non-TD *n* = 92, TD *n* = 120, Fig. 3a, Additional file 2: Table S6). The median absolute log_2_(fold change) was 0.12. The gene with the greatest fold change between Non-TD and TD subjects was TAS2R46 (log_2_(fold change) = 0.37, standard error = 0.12, Fig. 3b, Table 4). Further, the estimated fold changes of Non-TD-associated differentially-expressed genes were highly correlated between the individual studies (Pearson’s *r* = 0.80, *p* = 3.9E-16); however, fold changes of all transcripts were weakly correlated (Pearson’s r = 0.01, *p* = 0.10, Additional file 1: Fig. S7b). Additionally, median expression of differentially-expressed genes was not different from other genes on the array (MARBLES: differential = 4.48, non-differential = 4.64, *p* = 0.65; EARLI: differential = 4.15, non-differential = 4.20, *p* = 0.90; Additional file 1: Fig. S11).

**Figure 3.**
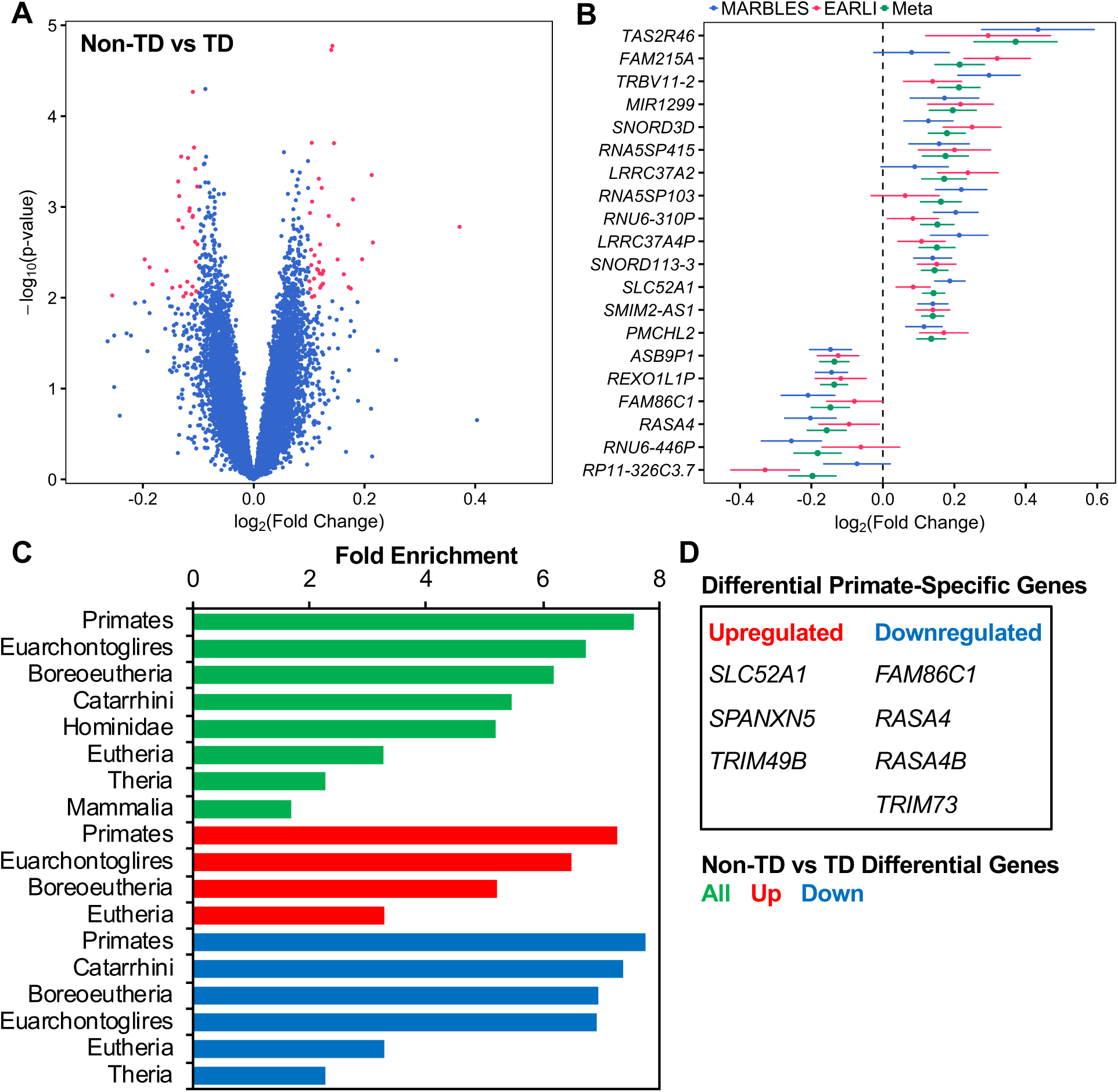
Identification and functional enrichment of Non-TD-associated differentially-expressed genes in cord blood. Gene expression in umbilical cord blood samples from subjects with typical development (TD, *n* = 120, 59 male/61 female) or those diagnosed with Non-TD at age 3 (*n* = 92, 50 male/42 female) was assessed by expression microarray. SVA was performed to control for technical and biological variables including sex and array batch. Differential expression analysis was carried out separately by study and combined in a meta-analysis. (A) Identification of 66 differentially-expressed genes in meta-analysis (66 probes, log_2_(fold change) > 0.1, p < 0.01). (B) Fold change in gene expression for top 20 differentially-expressed genes by meta-analysis log_2_(fold change) for individual studies and meta-analysis. (C) Non-TD-associated differentially-expressed genes were analyzed for functional enrichment in recently evolved genes with WebGestalt using the hypergeometric test. Significantly-enriched clades are shown (FDR *q* < 0.05). (D) Non-TD-associated differentially-expressed genes specific to primates are listed.

**Table 4.**
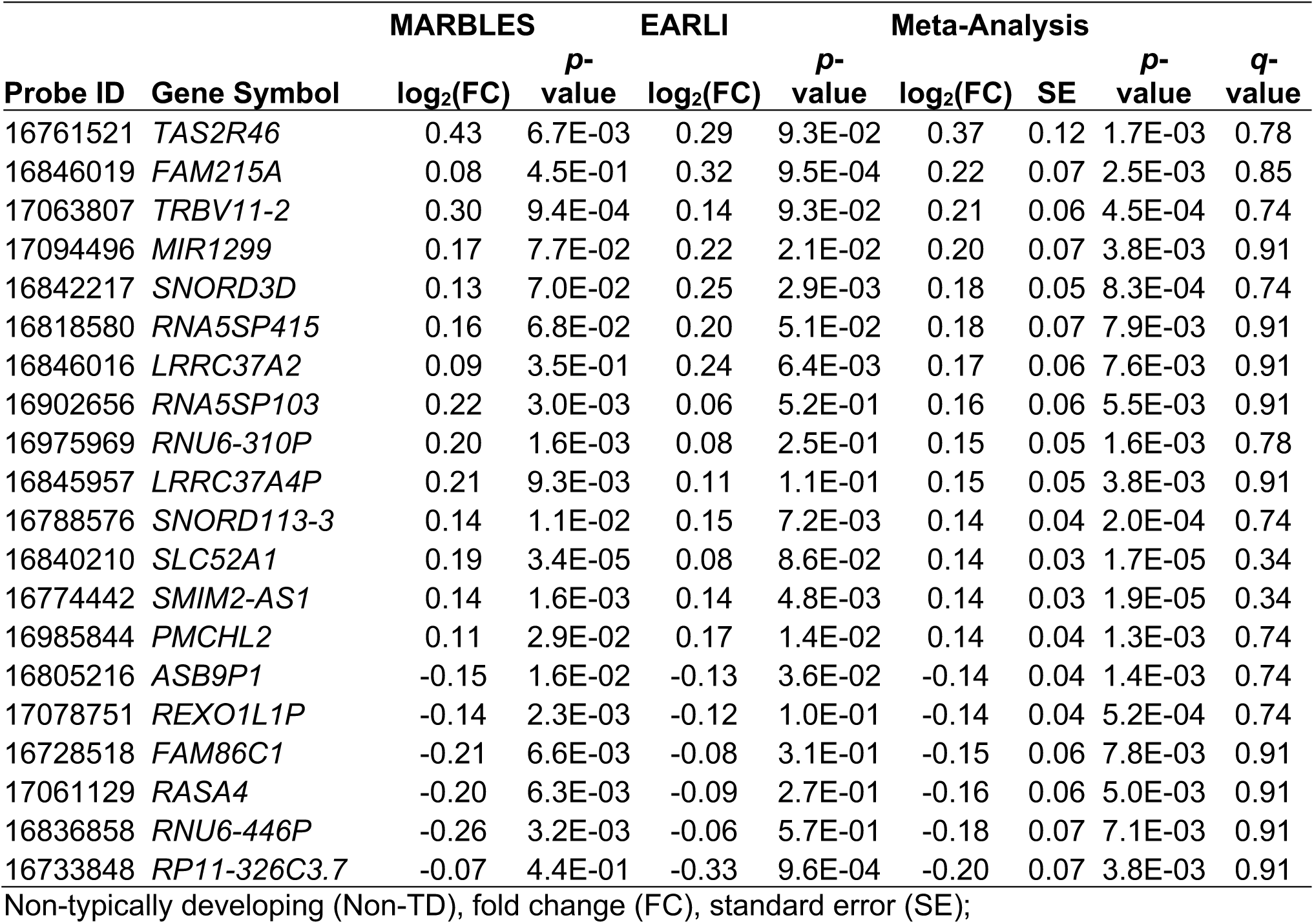
Top 20 Non-TD-associated differentially-expressed genes by log_2_(fold change) in meta-analysis.

Several of the 66 nominally differentially-expressed genes between Non-TD and TD cord blood samples, have been previously associated with genetic variation or gene expression in ASD, although the overlap was not statistically significant (*q* > 0.05, Additional file 1: Fig. S9, Additional file 2: Table S2). Genetic deficiencies in *MIR4269* have been previously associated with a reduced risk for ASD [43], while *DHCR24*, *GNAO1*, and *TYMS* were differentially expressed in ASD in other studies [8, 44]. Additionally, none of the Non-TD differentially-expressed genes were known cell-type specific genes (Additional file 1: Fig. S10, Additional file 2: Table S3)[41]. Differentially expressed genes in Non-TD that overlap genes previously associated with ASD likely function in general neurodevelopment.

Because genes recently evolved in primates have been hypothesized to play a role in human neurodevelopment, differentially-expressed genes in Non-TD cord blood were assessed for enrichment in recently evolved genes by vertebrate lineages, ranging from tetrapods to homininae using overrepresentation enrichment analysis [48]. Non-TD-associated genes were significantly enriched for genes recently evolved in mammalia, theria, eutheria, boreoeutheria, euarchontoglires, primates, catarrhini, and hominidae, with the greatest enrichment in primate-specific genes (fold enrichment = 7.5, *q* = 2.1E-5, Fig. 3c, Additional file 2: Table S7). Of genes that evolved in primates, *SLC52A1*, *SPANXN5*, and *TRIM49B* were upregulated in Non-TD cord blood, while *FAM86C1*, *RASA4*, *RASA4B*, and *TRIM73* were downregulated (Fig. 3d). In contrast, ASD differentially-expressed genes were not significantly enriched in recently evolved genes from any of the vertebrate lineages (*q* > 0.05).

After GSEA with all probes ranked by fold change in Non-TD compared to TD subjects, we observed significant enrichment for upregulation of sensory perception gene sets (*q* < 0.05, Fig. 4a, Additional file 2: Table S8). Taste receptor activity (14 LE transcripts, NES = 2.30, *q* < 1E-4), metabotropic glutamate receptors (17 LE transcripts, NES = 2.21, *q* = 4.9E-3), and olfactory receptor activity (105 LE transcripts, NES = 1.96, *q* = 0.018) gene sets were all upregulated in cord blood from Non-TD subjects. Additionally, gene sets that interact with the compounds quinine (19 LE transcripts, NES = 2.30, *q* = 2.0E-3) and citric acid (22 LE transcripts, NES = 2.17, *q* = 2.5E-3) were significantly upregulated, while those interacting with indomethacin (18 LE transcripts, NES = −2.02, *q* = 0.037) and H2-receptor antagonists (6 LE transcripts, NES = - 2.03, *q* = 0.047) were downregulated. Taste receptor genes included in these gene sets and the top Non-TD-upregulated gene, *TAS2R46*, are located at the 12p13.2 locus, which was also enriched (16 LE transcripts, NES = 2.11, *q* = 8.3E-3).

**Figure 4.**
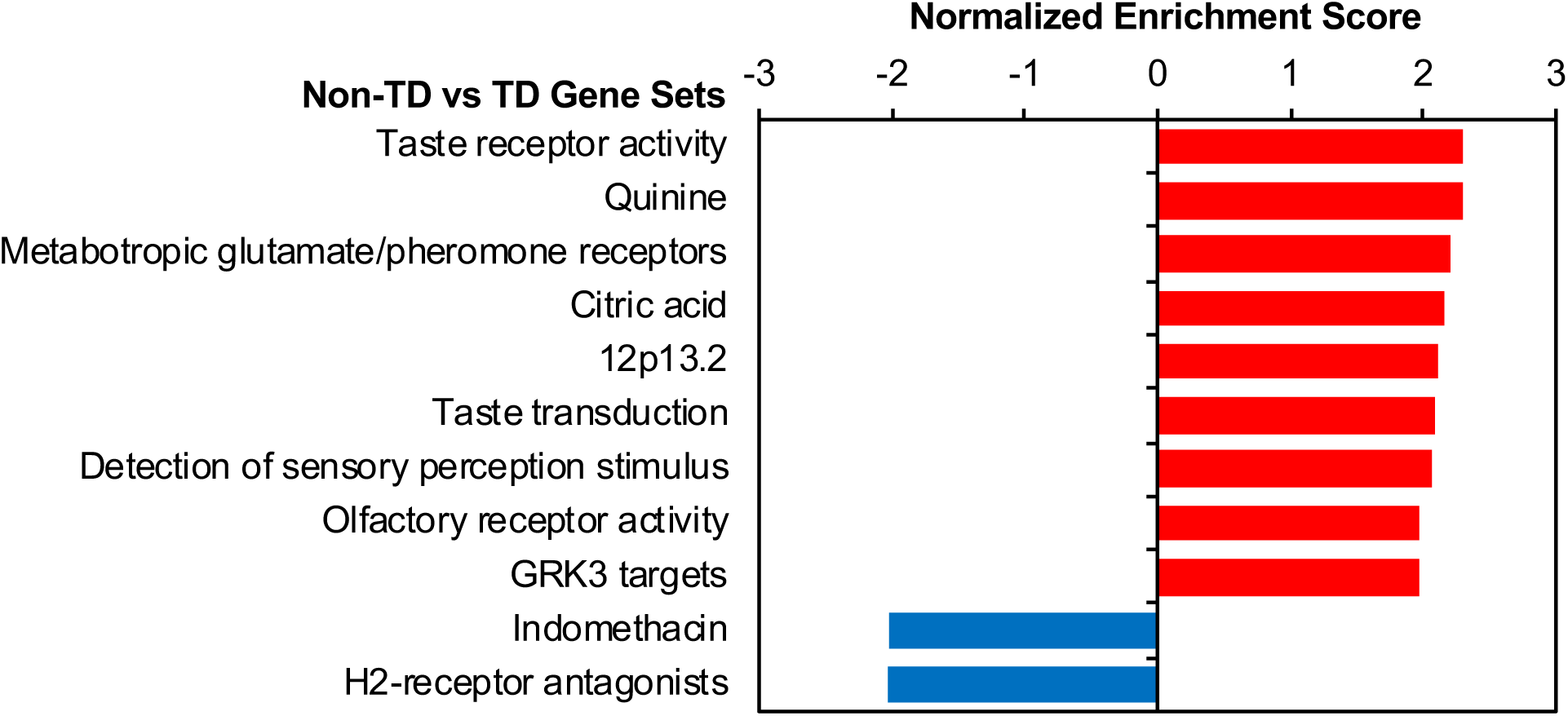
Sensory perception gene sets are dysregulated in Non-TD subject cord blood. Meta-analysis log_2_(fold change) for Non-TD versus TD gene expression was used to rank probes for GSEA with WebGestalt. GSEA assesses whether genes in biologically-defined sets occur toward the top or bottom of a ranked list more than expected by chance. Normalized enrichment score was plotted for significantly enriched gene sets (FDR *q* < 0.05) for default WebGestalt databases.

### Comparison of ASD and Non-TD differentially-expressed genes

Eight genes were differentially expressed in both ASD and Non-TD compared to TD subjects, which was more than expected by chance (odds ratio = 28.3, *p* = 1.67E-9, Fig. 5a). Specifically, *IGLV1-40*, *LRRC37A4P*, *MIR1299*, *PMCHL2*, and *TRBV11-2* were upregulated, while *RNU4ATAC11P*, *TPTE2P5, and TRIM74* were downregulated in both ASD and Non-TD subjects (Fig. 5b). *LRRC37AP*, *MIR1299*, *PMCHL2*, and *TRBV11-2* were among the top upregulated genes in Non-TD subjects (Fig. 3b). Additionally, the fold changes across all transcripts were moderately correlated between the ASD versus TD and Non-TD versus TD comparisons both within study and in the meta-analysis (Meta-analysis log_2_(fold change) Pearson’s r = 0.38, *p* < 2.2E-16, Additional file 1: Fig. S7c-e). These findings suggest that some ASD-associated transcriptional alterations in cord blood are also present in Non-TD subjects.

**Figure 5.**
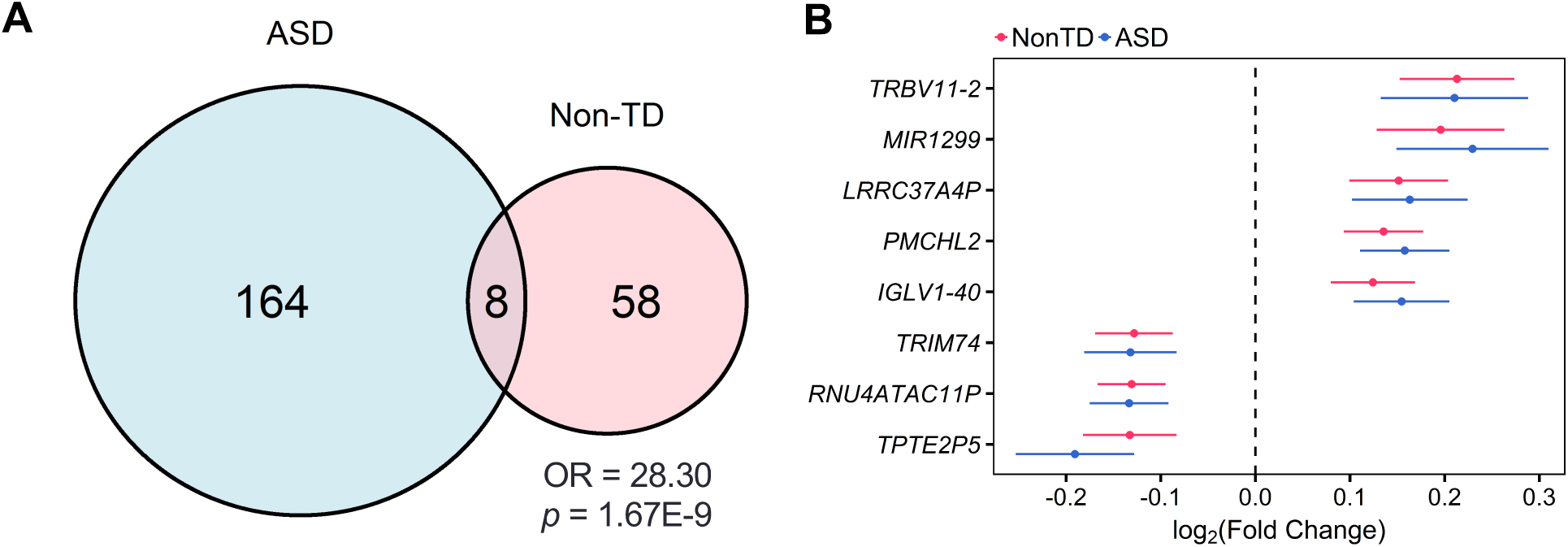
A subset of ASD-associated differentially-expressed genes are also differentially-expressed in Non-TD subjects. (A) Overlap of ASD- and Non-TD-associated differentially-expressed genes from meta-analysis by direction. Significance was assessed by Fisher’s Exact Test. (B) Meta-analysis log_2_(fold change) in gene expression for ASD- and Non-TD-associated differentially-expressed genes sorted by average log_2_(fold change).

### Coexpression network analysis and cell type deconvolution in cord blood from high-risk children

As a complementary approach to differential gene expression analysis, we performed WGCNA to identify consensus gene coexpression modules and their correlations with ASD or Non-TD diagnosis and other variables. 79 consensus coexpression modules were identified, which ranged in size from 20 to 4137 transcripts (Additional file 1: Fig. S12, Additional file 2: Table S9, S10). Overall the module eigengene networks were highly preserved between MARBLES and EARLI subjects, indicating the identified consensus modules are robust (Overall preservation = 0.93, Additional file 1: Fig. S13). Module eigengenes were correlated with diagnosis and demographic factors within each study and these results were combined in meta-analysis (Fig. 6, Additional file 1: Fig. S14-16, Additional file 2: Table S11). Across MARBLES and EARLI subjects, modules were identified that significantly correlated with diagnostic group, sex, gestational age, birthweight, ethnicity, paternal age, delivery method, and maternal smoking (FDR *q* < 0.05, Fig. 6, Additional file 1: Fig. S16). In particular, gestational age, birthweight, and paternal age were associated with more than 20 modules each, suggesting that these factors have major effects on gene expression in cord blood from high-risk children. Interestingly, the skyblue1 module was significantly upregulated in ASD compared to TD subjects (z-score = 3.4, FDR *q* = 0.046, Additional file 1: Fig. S17a-b, Additional file 2: Table S11). Skyblue1 was also significantly correlated with sex (z-score = −28.1, FDR *q* = 2.7E-172), paternal age (z-score = 2.2, FDR *q* = 0.047), and maternal smoking (z-score = −3.4, FDR *q* = 0.015). Probes in skyblue1 map to 21 genes, 18 of which are located only on chromosome Y, explaining the strong upregulation in males (Additional file 1: Fig. S17c). Further, two of these genes, *TTTY10* and *ZFY*, were upregulated in ASD compared to TD subjects in this study, which is a significant enrichment (Additional file 1: Fig. S17d, odds ratio = 33.6, *p* = 0.002). However, the association of skyblue1 with ASD may be driven by increased expression of skyblue1 genes in males and a greater frequency of males with ASD.

**Figure 6.**
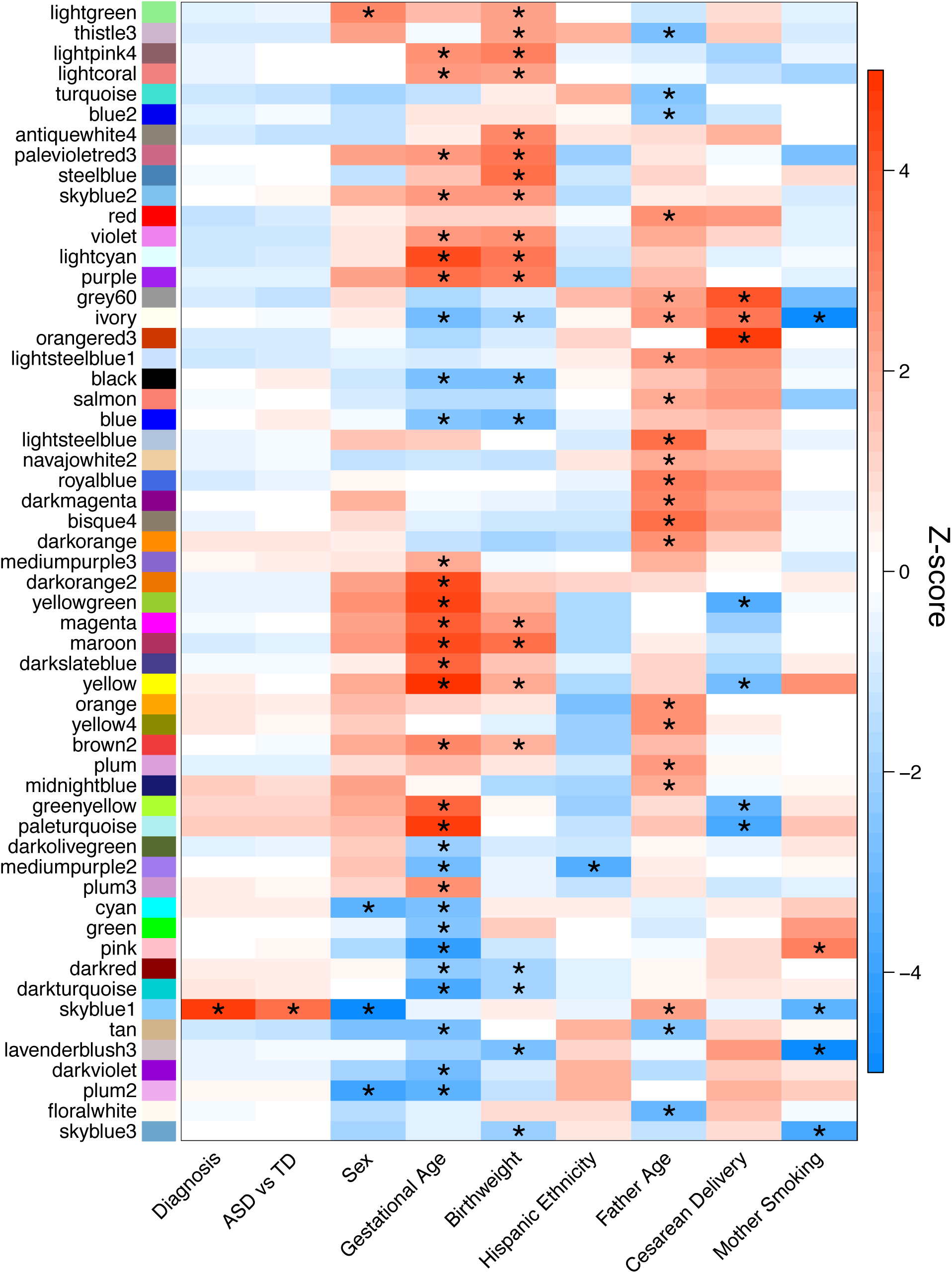
Consensus modules are correlated with diagnosis and demographic factors in meta-analysis. Heatmap of meta-analysis biweight midcorrelation Z-scores of module eigengenes with sample covariates (MARBLES: ASD *n* = 41, Non-TD *n* = 44, TD *n* = 76; EARLI: ASD *n* = 19, Non-TD *n* = 47, TD *n* = 43). Z-scores from the individual studies were combined using Stouffer’s method with weights given by the square root of sample *n.* P-values were adjusted for all comparisons shown in Additional file 1: Fig. S16 using the FDR method (* *q* < 0.05). Modules and covariates with significant correlations are shown.

Because cord blood is a heterogenous tissue composed of many cell types with distinct transcriptional profiles, we examined the proportions of 22 cell types estimated with the CIBERSORT web tool and their correlation with diagnostic group and demographic variables. The most prevalent cell types detected in cord blood overall were neutrophils and naïve CD4^+^ T cells, which made up 26% and 24% of whole cord blood, respectively (Additional file 2: Table S12). When cell type proportions were compared to diagnostic group and demographic factors within each study and combined in meta-analysis, no cell types differed between diagnostic groups (Fig. 7a). However, cell fractions were significantly correlated with demographic factors including sex, gestational age, birthweight, delivery method, and maternal smoking (FDR *q* < 0.05, Fig. 7b, Additional file 2: Table S13). To determine the correspondence of consensus modules with the proportions of specific cell types, the module eigengenes were compared to the cell type fractions followed by meta-analysis. The eigengenes for the majority of consensus modules were strongly and significantly correlated with the proportion of multiple cell types (FDR *q* < 0.05, Additional file 1: Fig. S18, Additional file 2: Table S14). The cell types significantly correlated with the most modules were neutrophils, naïve CD4^+^ T cells, CD8^+^ T cells, M0 macrophages, and plasma cells, which were associated with more than 30 modules each. In contrast, skyblue1, which was correlated with diagnosis, along with sex, paternal age, and maternal smoking, was not correlated with the proportions of any examined cell type (Additional file 1: Fig. S16, S18, Additional file 2: Table S14). As an example, the grey60 module was positively correlated with the proportions of naïve B cells, plasma cells, and activated dendritic cells, but negatively correlated with resting NK cells and neutrophils (FDR *q* < 0.05, Additional file 1: Fig. S18). The hub gene for grey60 was *CD79A*, which encodes Ig*α*, the B cell antigen receptor component necessary for signal transduction [53]. Interestingly, the grey60 module eigengene, *CD79A* expression, and the proportion of naïve B cells were all significantly upregulated during cesarean delivery (FDR *q* < 0.05) and nominally downregulated in ASD compared to Non-TD subjects (*p* < 0.05, Additional file 1: Fig. S19, Additional file 2: Table S11, S13). These results from coexpression and cell type deconvolution analyses suggest that biological factors including cell type, gestational age, birthweight, and paternal age are major drivers of interindividual variation in cord blood gene expression.

**Figure 7.**
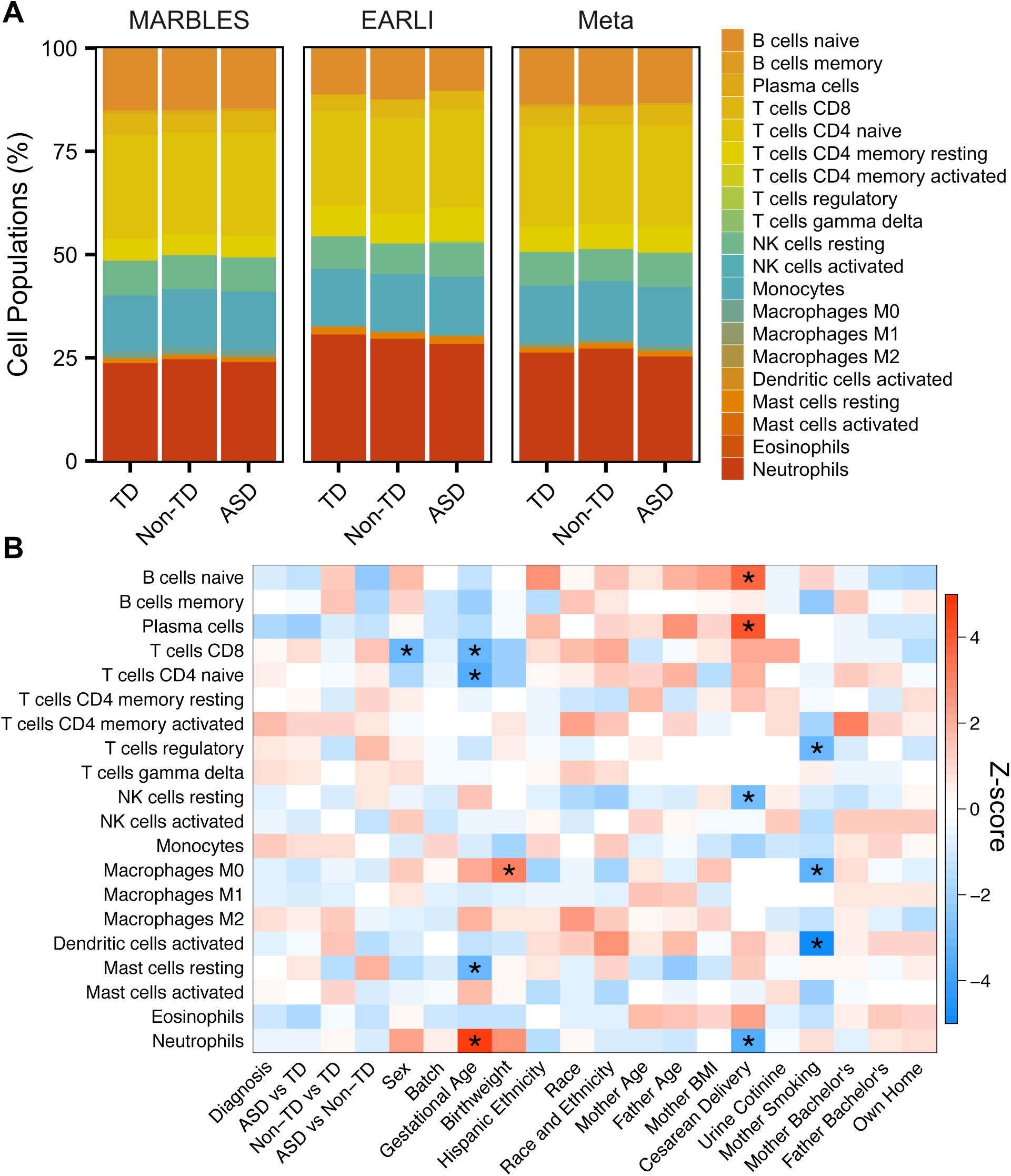
Cell type proportions are correlated with demographic factors in meta-analysis. (A) Barplot of mean estimated proportion of each cell type by diagnosis and study from CIBERSORT using default peripheral blood reference panel. (B) Heatmap of meta-analysis biweight midcorrelation Z-scores of cell type proportions with sample covariates (MARBLES: ASD *n* = 41, Non-TD *n* = 44, TD *n* = 76; EARLI: ASD *n* = 19, Non-TD *n* = 47, TD *n* = 43). Z-scores from the individual studies were combined using Stouffer’s method with weights given by the square root of sample *n.* P-values were adjusted using the FDR method (* *q* < 0.05).

## Discussion

### Perinatal transcriptional alterations in ASD

Based on meta-analysis across two high-risk pregnancy cohorts of similar design, we examined gene expression differences in cord blood between high-risk children who went on to develop ASD, were Non-TD, or were TD at 36 months. Significant differential gene expression in cord blood at individual genes was not observed in individuals who developed ASD compared to TD after adjusting for multiple comparisons. However, significant gene set enrichment was seen in toxic substance response, blood coagulation, chromatin regulation, and autoimmune response functions among genes differentially expressed at a nominal level in ASD.

Several of the nominally differentially-expressed genes have been previously associated with neurodevelopmental disorders such as ASD via genetic association studies. However, overlap with prior findings was not statistically significant, indicating that this may be coincidental. Included in the top upregulated genes in ASD was *TUBB2A*, a component of microtubules. *TUBB2A* is expressed in fetal brain and is mutated in complex cortical dysplasia, a neurodevelopmental disorder that involves seizures [54]. *SLC7A3* and *VSIG4* were also upregulated in ASD and have been previously associated as putative ASD risk genes [42]. Rare missense variants have been found in ASD subjects at *SLC7A3*, which is a cationic amino acid transporter selectively expressed in brain [55]. Nonsense and splice site variants have been found in ASD subjects at *VSIG4*, a complement C3 receptor that functions in pathogen recognition and inhibition of helper T-cell activation [56–58]. Overlap of both genetic and transcriptional association with ASD suggests that these genes would be interesting targets for additional mechanistic studies in relation to ASD.

Environmental factors are also thought to contribute to ASD risk, especially during the perinatal period, a sensitive window for neurodevelopment [59]. Genes differentially expressed in cord blood from ASD subjects, including *CYP1A1* and *GCH1*, were significantly enriched for functions in xenobiotic metabolism and response to both toxic substances and ultraviolet radiation. Notably, *CYP1A1* was downregulated in ASD cord blood and has been previously found to be transcriptionally regulated in blood by toxicants that affect neurodevelopment, including polychlorinated biphenyls [60–62]. *GCH1*--which is the rate-limiting enzyme in the synthesis of tetrahydrobiopterin, a precursor to folate, dopamine, and serotonin [63]--was also downregulated in cord blood from ASD subjects. *GCH1* expression increases in response to valproic acid, an anti-epileptic drug associated with behavioral deficits in mice and increased risk of autism in humans after *in utero* exposure [64, 65]. Interestingly, *GCH1* is genetically associated with ASD subphenotypes, is downregulated in peripheral blood from ASD children, and its product tetrahydrobiopterin is decreased in cerebrospinal fluid from ASD subjects [17, 66, 67]. These environmentally responsive genes may have altered expression due to increased genetic susceptibility and/or early life exposure to toxicants in patients with ASD [68, 69].

Epigenetic modifications, such as those to DNA and histone protein subunits, are affected by both genetic and environmental factors and are thought to play a role in mediating ASD risk [70, 71]. Immune dysregulation has also been found in children with ASD, and immune cells rely on epigenetic regulation for lineage commitment and cell activation in response to infection [72, 73]. A cluster of histone genes at 6p22.2 was significantly enriched for upregulated genes in ASD cord blood. Genes associated with the autoimmune disease systemic lupus erythematosus (SLE) were also upregulated, including histone-encoding, complement pathway, and antigen presentation genes. Epigenetic dysregulation is a feature of SLE, including global DNA hypomethylation, histone H3 and H4 hypoacetylation, and H3K9 hypomethylation in CD4+ T cells [74–76]. Notably, maternal SLE increases risk for ASD in offspring, suggesting an association between these two disorders [77]. Together, this implicates both epigenetic and immune mechanisms in ASD pathobiology.

### Perinatal transcriptional alterations in Non-TD

To assess the specificity of cord blood gene expression changes in ASD compared to other neurodevelopmental disorders, this analysis examined transcriptional differences in Non-TD compared to TD subjects across the two studies. While no single genes were significantly differentially expressed in Non-TD subjects after adjusting for multiple comparisons, sets of functionally-related genes were enriched among nominally differentially-expressed genes.

Significantly enriched gene sets included sensory perception and primate specific genes. The top upregulated gene in Non-TD cord blood was *TAS2R46*, encoding a member of the taste 2 receptor (TAS2R) family, which was included in the top upregulated gene set, taste receptor activity. Upregulated genes in this gene set were primarily other TAS2Rs located at the 12p13.2 locus. TAS2Rs are G-protein coupled receptors (GPCRs) highly expressed in taste receptor cells and are associated with the perception of bitter taste [78]. Interestingly, individuals with attention-deficit/hyperactivity disorder and epilepsy have previously been found to perceive bitter tastes more intensely than healthy controls [79, 80]. TAS2Rs are also expressed in blood leukocytes, where they function in chemosensation of foodborne flavor compounds and response to food uptake [81]. Further, TAS2Rs are upregulated in leukocytes from asthma patients and levels of lipopolysaccharide-induced pro-inflammatory cytokines are decreased by TAS2R agonists [82]. Taste receptor upregulation may reflect altered chemosensation in the immune and nervous systems in Non-TD subjects.

Differentially-expressed genes in cord blood from Non-TD subjects included genes recently evolved in primates and genes that function in neurodevelopment. Primate-specific genes originated at a similar evolutionary time that the neocortex expanded and have biased expression for the fetal neocortex in humans [48]. *RASA4* is a fetal-biased primate-specific gene that was also one of the top downregulated genes in Non-TD subjects. *RASA4* is a GTPase-activating protein in the Ras signaling pathway that functions in the activation of T cells, mast cells, and macrophages [83–85]. Children with Non-TD were also observed to have downregulation of *GNAO1*, encoding a G-protein alpha subunit important for neurodevelopment and synaptic signaling. Mutations in *GNAO1* are associated with epileptic encephalopathy, involuntary movements, and intellectual disability [86, 87]. Additionally, missense mutations and downregulation of *GNAO1* in lymphocytes occur in individuals with schizophrenia [88, 89]. In individuals with ASD, *GNAO1* is upregulated in post-mortem cortex [6]. Further, *GNAO1* is required in mast cells for toll-like receptor 2-mediated pro-inflammatory cytokine release, suggesting *GNAO1* functions in cell signaling in both the nervous and immune systems [90].

### Cord blood as a window into transcriptional alterations specific to ASD

Umbilical cord blood gene expression offers a unique snapshot of molecular differences in perinatal life, a critical window for neurodevelopment [91]. Hematopoietic cells in the blood are actively differentiating and responding to environmental cues, such as pathogens and xenobiotics [92, 93]. Epigenetic marks written during this period, which reflect short-term transcriptional activity, have the potential to have long-term effects on gene regulation and cell function [94, 95]. Signals from the immune system cross the blood-brain barrier during gestation and influence the development of the brain [96]. Toxicant exposure during gestation can also impact brain development [97, 98]. In this study, genes involved in toxic substance response, xenobiotic metabolism, and chromatin regulation were altered in cord blood from subjects diagnosed with ASD at 36 months. Transcriptional differences in cord blood from ASD and Non-TD subjects compared to TD subjects were largely independent, with only 8 genes in common. Enriched gene sets associated with Non-TD expression included sensory perception and primate-specific genes and did not overlap with ASD expression gene sets. Further, genes associated with ASD in previous studies of genetic variation and gene expression had few overlaps with ASD-associated genes in cord blood [6, 8, 42–44]. Instead, cord blood ASD genes likely represent tissue-specific environmentally-responsive genes that may reflect *in utero* exposures and long-term altered neurodevelopmental outcomes.

### Major factors contributing to transcriptional variability in whole cord blood

While the purpose of this study was to identify changes in cord blood gene expression associated with ASD and Non-TD diagnosis, we also identified biological factors associated with transcription variation in cord blood, including gestational age, birthweight, paternal age, and cell type composition. Gestational age was correlated with many consensus gene coexpression modules in cord blood, and it likely reflects ongoing differentiation and expansion of hematopoietic cells captured at different developmental timepoints. Specifically, gestational age was positively correlated with the estimated proportion of neutrophils and negatively correlated with the proportions of CD8^+^ T cells, naïve CD4^+^ T cells, and resting mast cells, suggesting alterations in these cell types are contributing to variability in gene expression related to gestational age. Notably, birthweight was correlated with many of the same gene coexpression modules as gestational age and is itself correlated with gestational age, so modules associated with birthweight likely reflect gestational age as well.

A mostly independent set of coexpression modules were correlated with paternal age. Increased paternal age has previously been associated with global changes in transcription in peripheral blood lymphocytes, including a downregulation of genes involved in transcriptional regulation and a decrease in the distribution of gene expression variance [99]. Interestingly, both increased paternal age and decreased gene expression variance have also been associated with ASD [99, 100]. The mechanism for the effect of paternal age on transcription is still unclear, but previous studies have observed increases in telomere length and *de novo* mutations and alterations in DNA methylation at specific genes in offspring from older fathers [101–103].

The factor with the strongest association with cord blood gene expression was cell type composition, as observed previously [104]. The cell types whose proportions correlated with the most modules were neutrophils, naïve CD4^+^ T cells, CD8^+^ T cells, M0 macrophages, and plasma cells. Neutrophils and naïve CD4^+^ T cells were also the most prevalent cell types in cord blood. The strong correlation of estimated cell type proportions with coexpression modules suggests that variability between samples in cell type composition is contributing a large portion of the transcriptional signal measured in whole cord blood. Overall, the large impact of biological factors on gene expression in a bulk tissue such as cord blood makes detecting differentially-expressed genes associated with a heterogeneous disorder such as ASD practically difficult. Future studies should take care to reduce these sources of noise, by isolating specific cell populations, selecting subjects with similar gestational and paternal age, or focusing on more narrow ASD endophenotypes [105].

## Limitations

Cord blood samples from two enriched autism risk pregnancy cohort studies were analyzed together to improve the power to detect differential gene expression between ASD and TD individuals. Meta-analysis substantially increased the total number of ASD and Non-TD subjects; however, the study was not adequately powered to detect moderate differences at a single gene level after correcting for multiple comparisons. Differential genes used in the enrichment analysis also did not meet multiple comparison-corrected statistical significance. Lack of statistical evidence of individual differential genes does not eliminate the potential of identifying a biologically-significant gene set enrichment across diagnostic groups; however, the results should be interpreted cautiously. ASD is a heterogeneous disorder and this may mask differential expression. The two cohort studies coordinated classifying participants into ASD, Non-TD, and TD diagnostic groups using the same metrics to improve consistency between the studies. Nonetheless, it is possible that heterogeneity within ASD remains in this study without further breaking down the group into subtypes.

The two cohorts used the same platform to measure the RNA expression levels and were subject to the limitations in transcript coverage and measurement precision of the microarray platform. Transcripts not covered by a probe on the array were not analyzed in this study, and targeted quantitative analysis in the general population would be needed to validate specific transcriptional changes as ASD risk biomarkers. Additionally, genetic, epigenetic, and environmental variation is known to impact both gene expression and ASD risk, but this was not investigated in this study. Future studies that integrate cord blood gene expression with genetic, epigenetic, and environmental factors will be important to improve understanding of ASD etiology.

## Conclusions

In the first study to investigate gene expression in cord blood from high-risk newborns later diagnosed with ASD, we identified nominally statistically significant transcriptional alterations specific to ASD, which were enriched for toxic substance response and epigenetic regulation functions. Differentially-expressed genes in ASD had few overlaps with those identified in cord blood from newborns with other non-typical neurodevelopmental outcomes in this high-risk population. Instead, Non-TD-associated genes were enriched for sensory perception functions and primate-specific genes. Further, coexpression and cell type analyses revealed that gestational age, birthweight, paternal age, and cell type composition have large impacts on variability in gene expression measured in whole cord blood.

A strength of this high-risk prospective pregnancy cohort design was the observation of gene expression at birth, prior to the onset of symptoms, diagnosis, and treatment. Perinatal life is a critical period for neurodevelopment, where environmental stressors could have long-term impact. Additionally, ASD-associated differential expression was meta-analyzed across two independent studies with harmonized methods and compared with expression changes in other subjects with non-typical neurodevelopment. Finally, cord blood is an accessible tissue that reflects the perinatal environment, and ASD-associated gene expression changes in cord blood may have potential as a predictive biomarker.

## List of abbreviations

Autism Diagnostic Interview-Revised (ADI-R), Autism Diagnostic Observation Schedule (ADOS), autism spectrum disorder (ASD), Early Autism Risk Longitudinal Investigation (EARLI), false discovery rate (FDR), fold change (FC), gene set enrichment analysis (GSEA), G-protein coupled receptors (GPCRs), histone deacetylase (HDAC), HUGO Gene Nomenclature Committee (HGNC), leading edge (LE), Markers of Autism Risk in Babies - Learning Early Signs (MARBLES), Mullen Scales of Early Learning (MSEL), natural killer cells (NK cells), non-typically developing (Non-TD), normalized enrichment score (NES), polycomb-repressive complex 2 (PRC2), robust multi-chip average (RMA), Simons Foundation Autism Research Initiative (SFARI), standard error (SE), surrogate variable analysis (SVA), systemic lupus erythematosus (SLE), taste 2 receptor (TAS2R), topological overlap matrix (TOM), typically developing (TD), University of California (UC), weighted gene correlation network analysis (WGCNA)

## Declarations

### Ethics approval and consent to participate

The UC Davis Institutional Review Board and the State of California Committee for the Protection of Human Subjects approved this study and the MARBLES Study protocols. Human Subjects Institutional Review Boards at each of the four sites in the EARLI Study approved this study and the EARLI Study protocols. Neither data nor specimens were collected until written informed consent was obtained from the parents.

### Consent for publication

Not applicable.

### Availability of data and material

The datasets supporting the conclusions of this article are available in the Gene Expression Omnibus (GEO) repository, under accession number GSE123302 at https://www.ncbi.nlm.nih.gov/geo/query/acc.cgi?acc=GSE123302. Data are shared if parents gave informed consent.

### Authors’ contributions

CEM and BYP were the lead authors and contributed substantially to data analysis, data interpretation, and drafting the manuscript. KMB contributed critical advice on data analysis methods and data interpretation. JIF and CL contributed to data analysis and interpretation. MDF, LAC, CJN, HEV, SO, and IH contributed to study design, as well as subject acquisition, diagnosis, and characterization. JML, RJS, and MDF conceived of the study and contributed substantially to data interpretation and manuscript revision. All authors read and approved the final manuscript.

## Acknowledgements

We would like to sincerely thank the participants in the MARBLES and EARLI studies.

## Additional Files

### Additional file 1.pdf

Supplemental Figure 1. Overview of study design.

Supplemental Figure 2. Surrogate variable analysis in MARBLES subjects.

Supplemental Figure 3. Surrogate variable analysis in EARLI subjects for the ASD versus TD comparison.

Supplemental Figure 4. Surrogate variable analysis in EARLI subjects for the Non-TD versus TD comparison.

Supplemental Figure 5. Identification of ASD-associated differentially-expressed genes in cord blood within each study.

Supplemental Figure 6. Identification of Non-TD-associated differentially-expressed genes in cord blood within each study.

Supplemental Figure 7. Correlations between ASD and Non-TD expression differences in MARBLES and EARLI subjects.

Supplemental Figure 8. Expression level distribution of meta-analysis ASD versus TD differential probes is similar to non-differential probes.

Supplemental Figure 9. Cord blood differentially-expressed genes are not enriched for ASD-associated gene sets.

Supplemental Figure 10. Cord blood differentially-expressed genes are depleted for blood cell-specific genes.

Supplemental Figure 11. Expression level distribution of meta-analysis Non-TD versus TD differential probes is similar to non-differential probes.

Supplemental Figure 12. Consensus coexpression modules identified in MARBLES and EARLI.

Supplemental Figure 13. Consensus module eigengene networks are preserved between

MARBLES and EARLI subjects.

Supplemental Figure 14. Consensus modules are correlated with diagnosis and demographic factors in MARBLES subjects.

Supplemental Figure 15. Consensus modules are correlated with demographic factors in EARLI subjects.

Supplemental Figure 16. Consensus modules are correlated with diagnosis and demographic factors in meta-analysis.

Supplemental Figure 17. Skyblue1 module is specifically expressed in males and is enriched for genes upregulated in ASD.

Supplemental Figure 18. Consensus modules are strongly correlated with cell type proportions in meta-analysis.

Supplemental Figure 19. The B cell-associated grey60 module is upregulated during cesarean delivery and is downregulated in ASD.

### Additional file 2.xlsx

Supplemental Table 1. Differential expression analysis of ASD compared to TD subjects for individual studies and meta-analysis.

Supplemental Table 2. Gene overlap analysis of ASD- and Non-TD-associated differentially-expressed genes with genes previously associated with ASD.

Supplemental Table 3. Gene overlap analysis of ASD- and Non-TD-associated differentially-expressed genes with blood cell-type specific genes.

Supplemental Table 4. Significant terms from overrepresentation enrichment analysis of ASD-associated differentially-expressed genes.

Supplemental Table 5. Significant terms from GSEA of probes ranked by ASD versus TD differential expression.

Supplemental Table 6. Differential expression analysis of Non-TD compared to TD subjects for individual studies and meta-analysis.

Supplemental Table 7. Significant terms from overrepresentation enrichment analysis of Non-TD-associated differentially-expressed genes.

Supplemental Table 8. Significant Terms from GSEA of probes ranked by Non-TD versus TD differential expression.

Supplemental Table 9. Consensus coexpression module membership in MARBLES subjects.

Supplemental Table 10. Consensus coexpression module membership in EARLI subjects.

Supplemental Table 11. Correlation meta-analysis of consensus module eigengenes with demographic factors.

Supplemental Table 12. Cell type proportions in MARBLES and EARLI subjects estimated with CIBERSORT.

Supplemental Table 13. Correlation meta-analysis of cell type proportions with demographic factors.

Supplemental Table 14. Correlation meta-analysis of consensus module eigengenes with cell type proportions.

## References

1. Constantino JN, Todorov A, Hilton C, Law P, Zhang Y, Molloy E, et al. Autism recurrence in half siblings: strong support for genetic mechanisms of transmission in ASD. Mol Psychiatry. 2012;18(2):137–8.

2. Sandin S, Lichtenstein P, Kuja-Halkola R, Larsson H, Hultman CM, Reichenberg A. The familial risk of autism. JAMA. 2014;311(17):1770–7.

3. Gaugler T, Klei L, Sanders SJ, Bodea CA, Goldberg AP, Lee AB, et al. Most genetic risk for autism resides with common variation. Nat Genet. 2014;46(8):881–5.

4. Weiner DJ, Wigdor EM, Ripke S, Walters RK, Kosmicki JA, Grove J, et al. Polygenic transmission disequilibrium confirms that common and rare variation act additively to create risk for autism spectrum disorders. Nat Genet. 2017;49(7):978–85.

5. Voineagu I, Wang X, Johnston P, Lowe JK, Tian Y, Horvath S, et al. Transcriptomic analysis of autistic brain reveals convergent molecular pathology. Nature. 2011;474(7351):380–4.

6. Gupta S, Ellis SE, Ashar FN, Moes A, Bader JS, Zhan J, et al. Transcriptome analysis reveals dysregulation of innate immune response genes and neuronal activity-dependent genes in autism. Nat Commun. 2014;5:5748.

7. Ansel A, Rosenzweig JP, Zisman PD, Melamed M, Gesundheit B. Variation in Gene Expression in Autism Spectrum Disorders: An Extensive Review of Transcriptomic Studies. Front Neurosci. 2016;10:601.

8. Tylee DS, Hess JL, Quinn TP, Barve R, Huang H, Zhang-James Y, et al. Blood transcriptomic comparison of individuals with and without autism spectrum disorder: A combined-samples mega-analysis. Am J Med Genet B Neuropsychiatr Genet. 2017;174(3):181–201.

9. Geschwind DH. Autism: many genes, common pathways? Cell. 2008;135(3):391–5.

10. De Rubeis S, He X, Goldberg AP, Poultney CS, Samocha K, Cicek AE, et al. Synaptic, transcriptional and chromatin genes disrupted in autism. Nature. 2014;515(7526):209–15.

11. Oblak AL, Rosene DL, Kemper TL, Bauman ML, Blatt GJ. Altered posterior cingulate cortical cyctoarchitecture, but normal density of neurons and interneurons in the posterior cingulate cortex and fusiform gyrus in autism. Autism Res. 2011;4(3):200–11.

12. Courchesne E, Pierce K. Brain overgrowth in autism during a critical time in development: implications for frontal pyramidal neuron and interneuron development and connectivity. Int J Dev Neurosci. 2005;23(2-3):153–70.

13. Bakulski KM, Feinberg JI, Andrews SV, Yang J, Brown S, S LM, et al. DNA methylation of cord blood cell types: Applications for mixed cell birth studies. Epigenetics. 2016;11(5):354–62.

14. Enstrom AM, Lit L, Onore CE, Gregg JP, Hansen RL, Pessah IN, et al. Altered gene expression and function of peripheral blood natural killer cells in children with autism. Brain Behav Immun. 2009;23(1):124–33.

15. Stamova B, Green PG, Tian Y, Hertz-Picciotto I, Pessah IN, Hansen R, et al. Correlations between gene expression and mercury levels in blood of boys with and without autism. Neurotox Res. 2011;19(1):31–48.

16. Tian Y, Green PG, Stamova B, Hertz-Picciotto I, Pessah IN, Hansen R, et al. Correlations of gene expression with blood lead levels in children with autism compared to typically developing controls. Neurotox Res. 2011;19(1):1–13.

17. Glatt SJ, Tsuang MT, Winn M, Chandler SD, Collins M, Lopez L, et al. Blood-based gene expression signatures of infants and toddlers with autism. J Am Acad Child Adolesc Psychiatry. 2012;51(9):934–44.e2.

18. Gregg JP, Lit L, Baron CA, Hertz-Picciotto I, Walker W, Davis RA, et al. Gene expression changes in children with autism. Genomics. 2008;91(1):22–9.

19. Kong SW, Collins CD, Shimizu-Motohashi Y, Holm IA, Campbell MG, Lee IH, et al. Characteristics and predictive value of blood transcriptome signature in males with autism spectrum disorders. PLoS One. 2012;7(12):e49475.

20. Kong SW, Shimizu-Motohashi Y, Campbell MG, Lee IH, Collins CD, Brewster SJ, et al. Peripheral blood gene expression signature differentiates children with autism from unaffected siblings. Neurogenetics. 2013;14(2):143–52.

21. Hertz-Picciotto I, Schmidt RJ, Walker CK, Bennett DH, Oliver M, Shedd-Wise KM, et al. A Prospective Study of Environmental Exposures and Early Biomarkers in Autism Spectrum Disorder: Design, Protocols, and Preliminary Data from the MARBLES Study. Environ Health Perspect. 2018;126(11):117004.

22. Newschaffer CJ, Croen LA, Fallin MD, Hertz-Picciotto I, Nguyen DV, Lee NL, et al. Infant siblings and the investigation of autism risk factors. J Neurodev Disord. 2012;4(1):7-1955–4-7.

23. Ozonoff S, Young GS, Carter A, Messinger D, Yirmiya N, Zwaigenbaum L, et al. Recurrence risk for autism spectrum disorders: a Baby Siblings Research Consortium study. Pediatrics. 2011;128(3):e488–95.

24. Lord C, Risi S, Lambrecht L, Cook EH, Jr., Leventhal BL, DiLavore PC, et al. The autism diagnostic observation schedule-generic: a standard measure of social and communication deficits associated with the spectrum of autism. J Autism Dev Disord. 2000;30(3):205–23.

25. Lord C, Rutter M, DiLavore PC, Risi S. The Autism Diagnostic Observation Schedule (ADOS). Los Angeles: Western Psychological Services; 2000.

26. Lord C, Rutter M, Le Couteur A. Autism Diagnostic Interview-Revised: a revised version of a diagnostic interview for caregivers of individuals with possible pervasive developmental disorders. J Autism Dev Disord. 1994;24(5):659–85.

27. Mullen EM. Scales of Early Learning. Circle Pines, MN: American Guidance Services Inc; 1995.

28. Ozonoff S, Young GS, Belding A, Hill M, Hill A, Hutman T, et al. The broader autism phenotype in infancy: when does it emerge? J Am Acad Child Adolesc Psychiatry. 2014;53(4):398-407 e2.

29. Chawarska K, Shic F, Macari S, Campbell DJ, Brian J, Landa R, et al. 18-month predictors of later outcomes in younger siblings of children with autism spectrum disorder: a baby siblings research consortium study. J Am Acad Child Adolesc Psychiatry. 2014;53(12):1317–27 e1.

30. Verification SSoB. Biochemical verification of tobacco use and cessation. Nicotine Tob Res. 2002;4(2):149–59.

31. Irizarry RA, Hobbs B, Collin F, Beazer-Barclay YD, Antonellis KJ, Scherf U, et al. Exploration, normalization, and summaries of high density oligonucleotide array probe level data. Biostatistics. 2003;4(2):249–64.

32. Kauffmann A, Gentleman R, Huber W. arrayQualityMetrics--a bioconductor package for quality assessment of microarray data. Bioinformatics. 2009;25(3):415–6.

33. Carvalho BS, Irizarry RA. A framework for oligonucleotide microarray preprocessing. Bioinformatics. 2010;26(19):2363–7.

34. Carvalho B. pd.hugene.2.0.st: Platform Design Info for Affymetrix HuGene-2_0-st. R package. 2015;version 3.14.1:https://bioconductor.org/packages/release/data/annotation/html/pd.hugene.2.0.st.html.

35. Leek JT, Storey JD. Capturing heterogeneity in gene expression studies by surrogate variable analysis. PLoS Genet. 2007;3(9):1724–35.

36. Leek JT, Johnson WE, Parker HS, Jaffe AE, Storey JD. The sva package for removing batch effects and other unwanted variation in high-throughput experiments. Bioinformatics. 2012;28(6):882–3.

37. Hoffman GE, Schadt EE. variancePartition: interpreting drivers of variation in complex gene expression studies. BMC Bioinformatics. 2016;17(1):483.

38. Ritchie ME, Phipson B, Wu D, Hu Y, Law CW, Shi W, et al. limma powers differential expression analyses for RNA-sequencing and microarray studies. Nucleic Acids Res. 2015;43(7):e47.

39. Willer CJ, Li Y, Abecasis GR. METAL: fast and efficient meta-analysis of genomewide association scans. Bioinformatics. 2010;26(17):2190–1.

40. Shen L, Sinai M. GeneOverlap: Test and visualize gene overlaps. R package. 2013; version 1.16.0:http://shenlab-sinai.github.io/shenlab-sinai/.

41. Newman AM, Liu CL, Green MR, Gentles AJ, Feng W, Xu Y, et al. Robust enumeration of cell subsets from tissue expression profiles. Nat Methods. 2015;12(5):453–7.

42. Abrahams BS, Arking DE, Campbell DB, Mefford HC, Morrow EM, Weiss LA, et al. SFARI Gene 2.0: a community-driven knowledgebase for the autism spectrum disorders (ASDs). Mol Autism. 2013;4(1):36.

43. Autism Spectrum Disorders Working Group of The Psychiatric Genomics Consortium. Meta-analysis of GWAS of over 16,000 individuals with autism spectrum disorder highlights a novel locus at 10q24.32 and a significant overlap with schizophrenia. Mol Autism. 2017;8:21.

44. Parikshak NN, Swarup V, Belgard TG, Irimia M, Ramaswami G, Gandal MJ, et al. Genome-wide changes in lncRNA, splicing, and regional gene expression patterns in autism. Nature. 2016;540(7633):423–7.

45. Tylee DS, Espinoza AJ, Hess JL, Tahir MA, McCoy SY, Rim JK, et al. RNA sequencing of transformed lymphoblastoid cells from siblings discordant for autism spectrum disorders reveals transcriptomic and functional alterations: Evidence for sex-specific effects. Autism Res. 2017;10(3):439–55.

46. Durinck S, Spellman PT, Birney E, Huber W. Mapping identifiers for the integration of genomic datasets with the R/Bioconductor package biomaRt. Nat Protoc. 2009;4(8):1184–91.

47. Wang J, Duncan D, Shi Z, Zhang B. WEB-based GEne SeT AnaLysis Toolkit (WebGestalt): update 2013. Nucleic Acids Res. 2013;41(Web Server issue):W77–83.

48. Zhang YE, Landback P, Vibranovski MD, Long M. Accelerated recruitment of new brain development genes into the human genome. PLoS Biol. 2011;9(10):e1001179.

49. Subramanian A, Tamayo P, Mootha VK, Mukherjee S, Ebert BL, Gillette MA, et al. Gene set enrichment analysis: a knowledge-based approach for interpreting genome-wide expression profiles. Proc Natl Acad Sci U S A. 2005;102(43):15545–50.

50. Langfelder P, Horvath S. WGCNA: an R package for weighted correlation network analysis. BMC Bioinformatics. 2008;9:559.

51. Langfelder P, Mischel PS, Horvath S. When is hub gene selection better than standard meta-analysis? PLoS One. 2013;8(4):e61505.

52. Consortium ASDWGoTPG. Meta-analysis of GWAS of over 16,000 individuals with autism spectrum disorder highlights a novel locus at 10q24.32 and a significant overlap with schizophrenia. Mol Autism. 2017;8:21.

53. Wienands J, Engels N. Multitasking of Ig-alpha and Ig-beta to regulate B cell antigen receptor function. Int Rev Immunol. 2001;20(6):679–96.

54. Cushion TD, Paciorkowski AR, Pilz DT, Mullins JG, Seltzer LE, Marion RW, et al. De novo mutations in the beta-tubulin gene TUBB2A cause simplified gyral patterning and infantile-onset epilepsy. Am J Hum Genet. 2014;94(4):634–41.

55. Nava C, Rupp J, Boissel JP, Mignot C, Rastetter A, Amiet C, et al. Hypomorphic variants of cationic amino acid transporter 3 in males with autism spectrum disorders. Amino Acids. 2015;47(12):2647–58.

56. Lim ET, Raychaudhuri S, Sanders SJ, Stevens C, Sabo A, MacArthur DG, et al. Rare complete knockouts in humans: population distribution and significant role in autism spectrum disorders. Neuron. 2013;77(2):235–42.

57. van Lookeren Campagne M, Verschoor A. Pathogen clearance and immune adherence “revisited”: Immuno-regulatory roles for CRIg. Semin Immunol. 2018;37:4–11.

58. Jung K, Seo SK, Choi I. Endogenous VSIG4 negatively regulates the helper T cell-mediated antibody response. Immunol Lett. 2015;165(2):78–83.

59. Kalkbrenner AE, Daniels JL, Chen JC, Poole C, Emch M, Morrissey J. Perinatal exposure to hazardous air pollutants and autism spectrum disorders at age 8. Epidemiology. 2010;21(5):631–41.

60. Leijs MM, Esser A, Amann PM, Schettgen T, Heise R, Fietkau K, et al. Expression of CYP1A1, CYP1B1 and IL-1beta in PBMCs and skin samples of PCB exposed individuals. Sci Total Environ. 2018;642:1429–38.

61. Vorrink SU, Hudachek DR, Domann FE. Epigenetic determinants of CYP1A1 induction by the aryl hydrocarbon receptor agonist 3,3’,4,4’,5-pentachlorobiphenyl (PCB 126). Int J Mol Sci. 2014;15(8):13916–31.

62. Park HY, Hertz-Picciotto I, Sovcikova E, Kocan A, Drobna B, Trnovec T. Neurodevelopmental toxicity of prenatal polychlorinated biphenyls (PCBs) by chemical structure and activity: a birth cohort study. Environ Health. 2010;9:51.

63. Thony B, Auerbach G, Blau N. Tetrahydrobiopterin biosynthesis, regeneration and functions. Biochem J. 2000;347 Pt 1:1–16.

64. Colleoni S, Galli C, Gaspar JA, Meganathan K, Jagtap S, Hescheler J, et al. A comparative transcriptomic study on the effects of valproic acid on two different hESCs lines in a neural teratogenicity test system. Toxicol Lett. 2014;231(1):38–44.

65. Roullet FI, Lai JK, Foster JA. In utero exposure to valproic acid and autism--a current review of clinical and animal studies. Neurotoxicol Teratol. 2013;36:47–56.

66. Hu VW, Addington A, Hyman A. Novel autism subtype-dependent genetic variants are revealed by quantitative trait and subphenotype association analyses of published GWAS data. PLoS One. 2011;6(4):e19067.

67. Tani Y, Fernell E, Watanabe Y, Kanai T, Langstrom B. Decrease in 6R-5,6,7,8-tetrahydrobiopterin content in cerebrospinal fluid of autistic patients. Neurosci Lett. 1994;181(1-2):169–72.

68. Herbert MR, Russo JP, Yang S, Roohi J, Blaxill M, Kahler SG, et al. Autism and environmental genomics. Neurotoxicology. 2006;27(5):671–84.

69. Kalkbrenner AE, Schmidt RJ, Penlesky AC. Environmental chemical exposures and autism spectrum disorders: a review of the epidemiological evidence. Curr Probl Pediatr Adolesc Health Care. 2014;44(10):277–318.

70. LaSalle JM. Epigenomic strategies at the interface of genetic and environmental risk factors for autism. J Hum Genet. 2013;58(7):396–401.

71. Tordjman S, Somogyi E, Coulon N, Kermarrec S, Cohen D, Bronsard G, et al. Gene x Environment interactions in autism spectrum disorders: role of epigenetic mechanisms. Front Psychiatry. 2014;5:53.

72. Zhao M, Wang Z, Yung S, Lu Q. Epigenetic dynamics in immunity and autoimmunity. Int J Biochem Cell Biol. 2015;67:65–74.

73. Ashwood P, Krakowiak P, Hertz-Picciotto I, Hansen R, Pessah I, Van de Water J. Elevated plasma cytokines in autism spectrum disorders provide evidence of immune dysfunction and are associated with impaired behavioral outcome. Brain Behav Immun. 2011;25(1):40–5.

74. Long H, Yin H, Wang L, Gershwin ME, Lu Q. The critical role of epigenetics in systemic lupus erythematosus and autoimmunity. J Autoimmun. 2016;74:118–38.

75. Coit P, Jeffries M, Altorok N, Dozmorov MG, Koelsch KA, Wren JD, et al. Genome-wide DNA methylation study suggests epigenetic accessibility and transcriptional poising of interferon-regulated genes in naive CD4+ T cells from lupus patients. J Autoimmun. 2013;43:78–84.

76. Hu N, Qiu X, Luo Y, Yuan J, Li Y, Lei W, et al. Abnormal histone modification patterns in lupus CD4+ T cells. J Rheumatol. 2008;35(5):804–10.

77. Vinet E, Pineau CA, Clarke AE, Scott S, Fombonne E, Joseph L, et al. Increased Risk of Autism Spectrum Disorders in Children Born to Women With Systemic Lupus Erythematosus: Results From a Large Population-Based Cohort. Arthritis Rheumatol. 2015;67(12):3201–8.

78. Conte C, Ebeling M, Marcuz A, Nef P, Andres-Barquin PJ. Identification and characterization of human taste receptor genes belonging to the TAS2R family. Cytogenet Genome Res. 2002;98(1):45–53.

79. Weiland R, Macht M, Ellgring H, Gross-Lesch S, Lesch KP, Pauli P. Olfactory and gustatory sensitivity in adults with attention-deficit/hyperactivity disorder. Atten Defic Hyperact Disord. 2011;3(1):53–60.

80. Campanella G, Filla A, De Michele G. Smell and taste acuity in epileptic syndromes. Eur Neurol. 1978;17(3):136–41.

81. Malki A, Fiedler J, Fricke K, Ballweg I, Pfaffl MW, Krautwurst D. Class I odorant receptors, TAS1R and TAS2R taste receptors, are markers for subpopulations of circulating leukocytes. J Leukoc Biol. 2015;97(3):533–45.

82. Orsmark-Pietras C, James A, Konradsen JR, Nordlund B, Soderhall C, Pulkkinen V, et al. Transcriptome analysis reveals upregulation of bitter taste receptors in severe asthmatics. Eur Respir J. 2013;42(1):65–78.

83. Strazza M, Azoulay-Alfaguter I, Dun B, Baquero-Buitrago J, Mor A. CD28 inhibits T cell adhesion by recruiting CAPRI to the plasma membrane. J Immunol. 2015;194(6):2871–7.

84. Nakamura R, Furuno T, Nakanishi M. The plasma membrane shuttling of CAPRI is related to regulation of mast cell activation. Biochem Biophys Res Commun. 2006;347(1):363–8.

85. Zhang J, Guo J, Dzhagalov I, He YW. An essential function for the calcium-promoted Ras inactivator in Fcgamma receptor-mediated phagocytosis. Nat Immunol. 2005;6(9):911–9.

86. Nakamura K, Kodera H, Akita T, Shiina M, Kato M, Hoshino H, et al. De Novo mutations in GNAO1, encoding a Galphao subunit of heterotrimeric G proteins, cause epileptic encephalopathy. Am J Hum Genet. 2013;93(3):496–505.

87. Saitsu H, Fukai R, Ben-Zeev B, Sakai Y, Mimaki M, Okamoto N, et al. Phenotypic spectrum of GNAO1 variants: epileptic encephalopathy to involuntary movements with severe developmental delay. Eur J Hum Genet. 2016;24(1):129–34.

88. Vawter MP, Ferran E, Galke B, Cooper K, Bunney WE, Byerley W. Microarray screening of lymphocyte gene expression differences in a multiplex schizophrenia pedigree. Schizophr Res. 2004;67(1):41–52.

89. Tani M, Mui K, Minami Y, Kiriike N. Association of a GTP-binding protein Go alpha subunit mutation with schizophrenia. Mol Psychiatry. 2001;6(4):359.

90. Jin M, Yu B, Zhang W, Zhang W, Xiao Z, Mao Z, et al. Toll-like receptor 2-mediated MAPKs and NF-kappaB activation requires the GNAO1-dependent pathway in human mast cells. Integr Biol (Camb). 2016;8(9):968–75.

91. Richetto J, Riva MA. Prenatal maternal factors in the development of cognitive impairments in the offspring. J Reprod Immunol. 2014;104–105:20-5.

92. Lura MP, Gorlanova O, Muller L, Proietti E, Vienneau D, Reppucci D, et al. Response of cord blood cells to environmental, hereditary and perinatal factors: A prospective birth cohort study. PLoS One. 2018;13(7):e0200236.

93. Cardenas A, Koestler DC, Houseman EA, Jackson BP, Kile ML, Karagas MR, et al. Differential DNA methylation in umbilical cord blood of infants exposed to mercury and arsenic in utero. Epigenetics. 2015;10(6):508–15.

94. Vineis P, Chatziioannou A, Cunliffe VT, Flanagan JM, Hanson M, Kirsch-Volders M, et al. Epigenetic memory in response to environmental stressors. Faseb j. 2017;31(6):2241–51.

95. Richmond RC, Simpkin AJ, Woodward G, Gaunt TR, Lyttleton O, McArdle WL, et al. Prenatal exposure to maternal smoking and offspring DNA methylation across the lifecourse: findings from the Avon Longitudinal Study of Parents and Children (ALSPAC). Hum Mol Genet. 2015;24(8):2201–17.

96. Bilbo SD, Schwarz JM. The immune system and developmental programming of brain and behavior. Front Neuroendocrinol. 2012;33(3):267–86.

97. Lin CC, Chien CJ, Tsai MS, Hsieh CJ, Hsieh WS, Chen PC. Prenatal phenolic compounds exposure and neurobehavioral development at 2 and 7years of age. Sci Total Environ. 2017;605–606:801-10.

98. Perera F, Phillips DH, Wang Y, Roen E, Herbstman J, Rauh V, et al. Prenatal exposure to polycyclic aromatic hydrocarbons/aromatics, BDNF and child development. Environ Res. 2015;142:602–8.

99. Alter MD, Kharkar R, Ramsey KE, Craig DW, Melmed RD, Grebe TA, et al. Autism and increased paternal age related changes in global levels of gene expression regulation. PLoS One. 2011;6(2):e16715.

100. Hultman CM, Sandin S, Levine SZ, Lichtenstein P, Reichenberg A. Advancing paternal age and risk of autism: new evidence from a population-based study and a meta-analysis of epidemiological studies. Mol Psychiatry. 2011;16(12):1203–12.

101. Atsem S, Reichenbach J, Potabattula R, Dittrich M, Nava C, Depienne C, et al. Paternal age effects on sperm FOXK1 and KCNA7 methylation and transmission into the next generation. Hum Mol Genet. 2016;25(22):4996–5005.

102. Unryn BM, Cook LS, Riabowol KT. Paternal age is positively linked to telomere length of children. Aging Cell. 2005;4(2):97–101.

103. Girard SL, Bourassa CV, Lemieux Perreault LP, Legault MA, Barhdadi A, Ambalavanan A, et al. Paternal Age Explains a Major Portion of De Novo Germline Mutation Rate Variability in Healthy Individuals. PLoS One. 2016;11(10):e0164212.

104. Schurmann C, Heim K, Schillert A, Blankenberg S, Carstensen M, Dorr M, et al. Analyzing illumina gene expression microarray data from different tissues: methodological aspects of data analysis in the metaxpress consortium. PLoS One. 2012;7(12):e50938.

105. Constantino JN. Deconstructing autism: from unitary syndrome to contributory developmental endophenotypes. Int Rev Psychiatry. 2018;30(1):18–24.

