## Additional File 1 for "A meta-analysis of two high-risk prospective cohort studies reveals autism-specific transcriptional changes to chromatin, autoimmune, and environmental response genes in umbilical cord blood"

**Supplemental Figures**

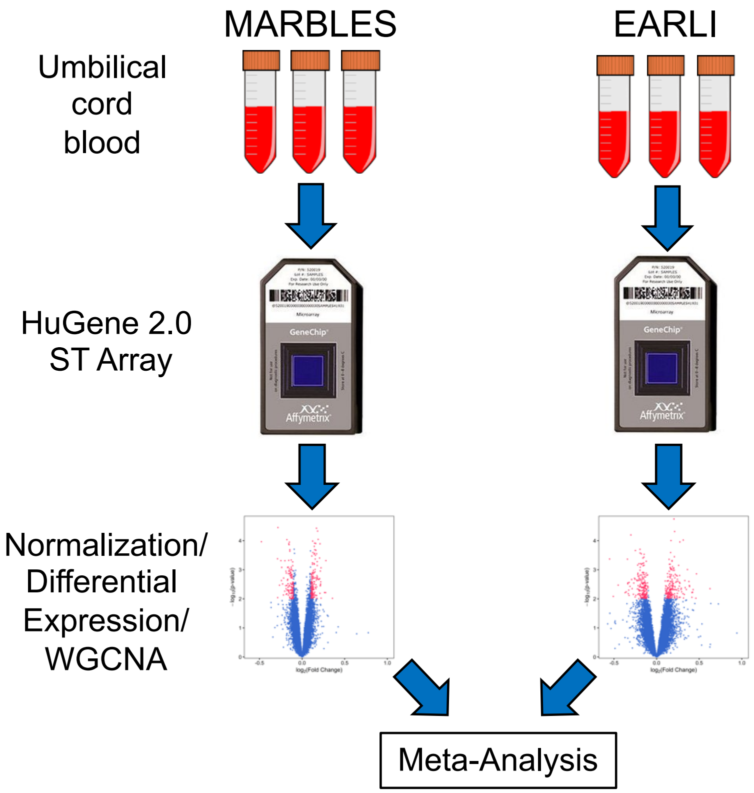


**Supplemental Figure 1. Overview of study design.** Umbilical cord blood was collected independently from participants in the high-risk prospective MARBLES and EARLI studies and frozen. After behavioral assessment at 36 months, RNA was extracted and assessed for gene expression with the HuGene 2.0 ST Array (Affymetrix). Probe intensities were normalized and analyzed for differential expression within each study. Differential expression analysis included SVA to control for technical and biological variables including array batch and sex. Probe fold change and standard error were combined across studies in the meta-analysis. Expression data was adjusted for batch before WGCNA. Consensus modules were identified and correlation z-scores were combined across studies in the meta-analysis.

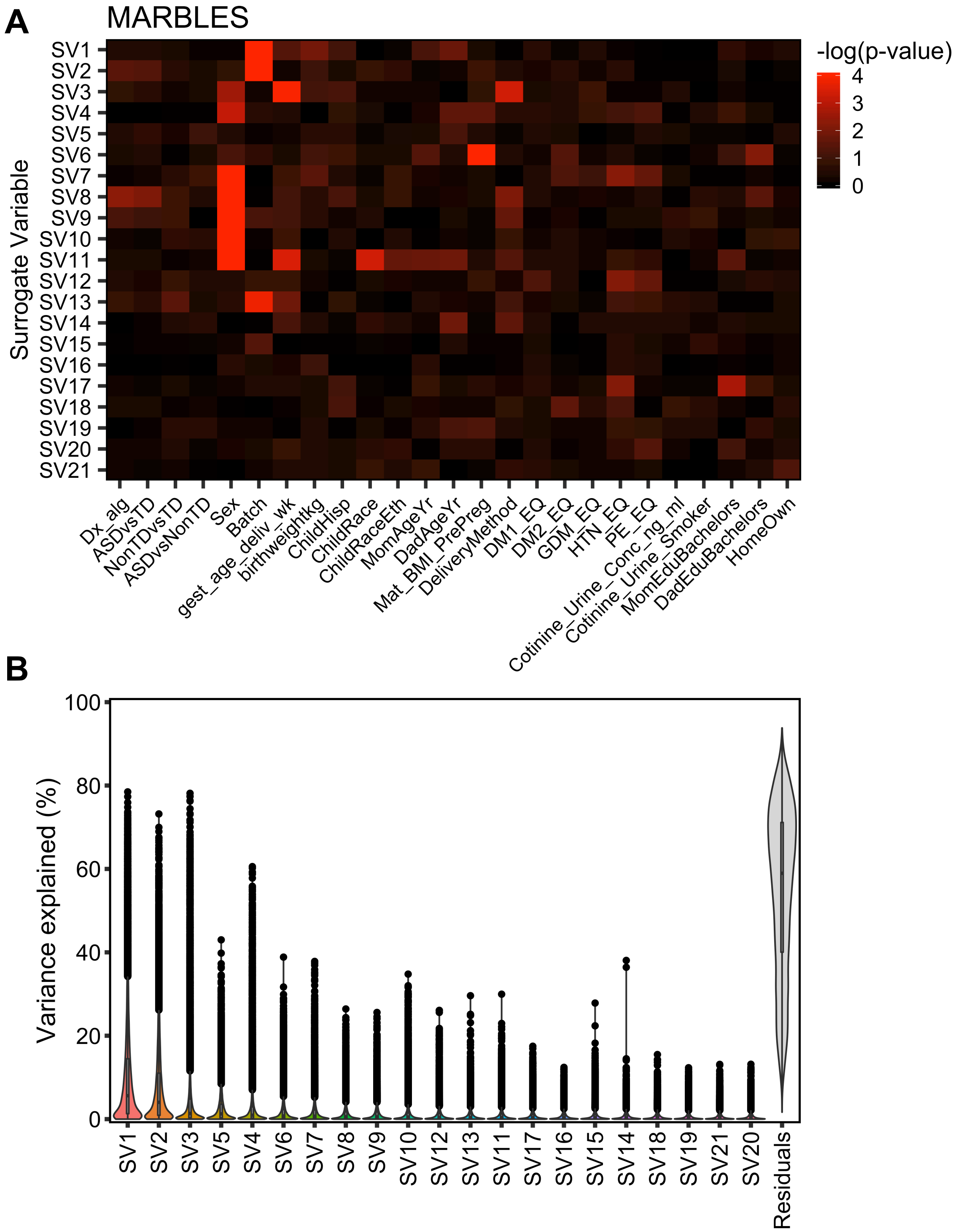


**Supplemental Figure 2. Surrogate variable analysis in MARBLES subjects.**

(A) Association of surrogate variables with covariates using linear regression. (B) Proportion of variance in expression of each probe explained by each surrogate variable, sorted by median variance explained.

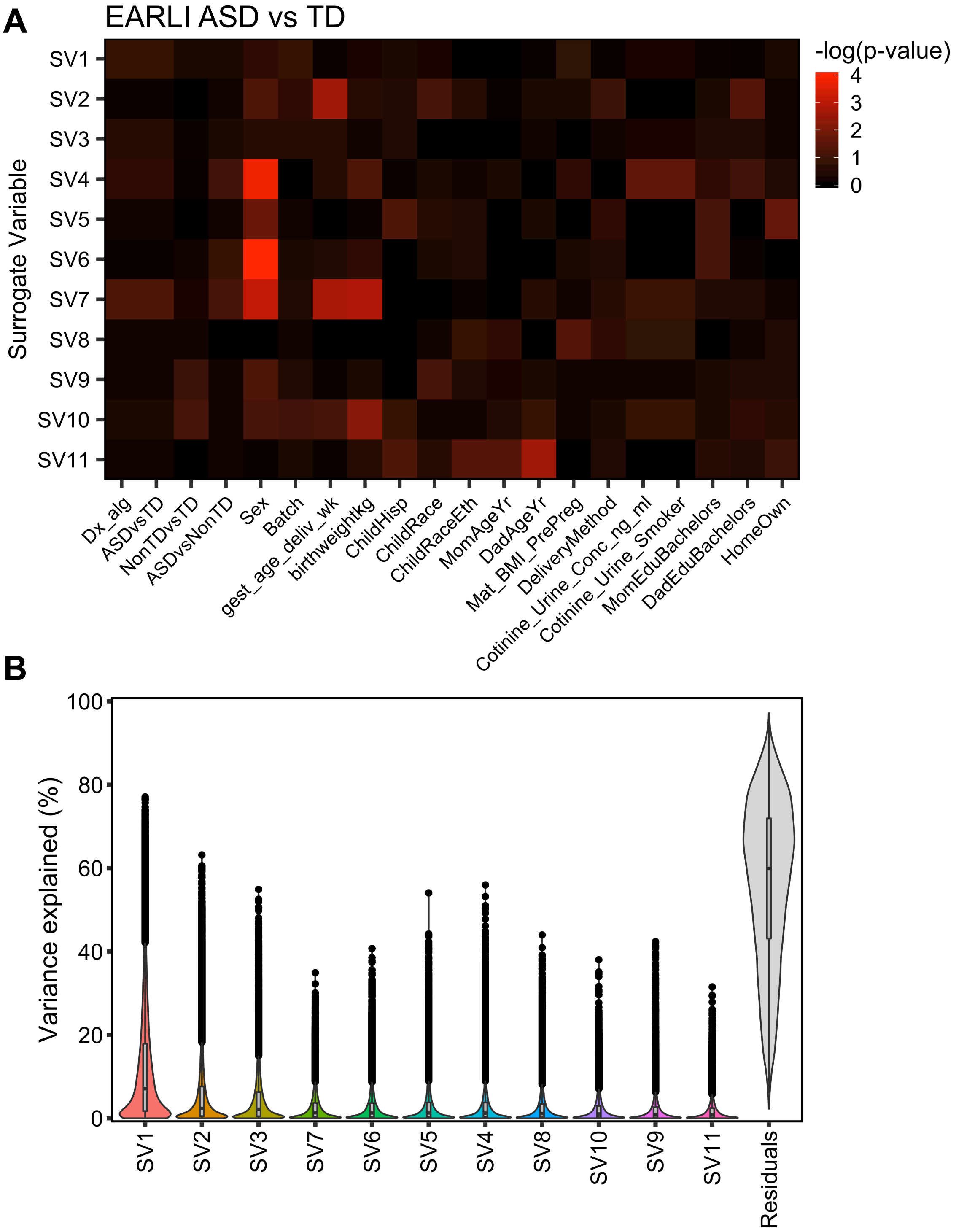

**Supplemental Figure 3. Surrogate variable analysis in EARLI subjects for the ASD versus TD comparison.** (A) Association of surrogate variables with covariates using linear regression. (B) Proportion of variance in expression of each probe explained by each surrogate variable, sorted by median variance explained.

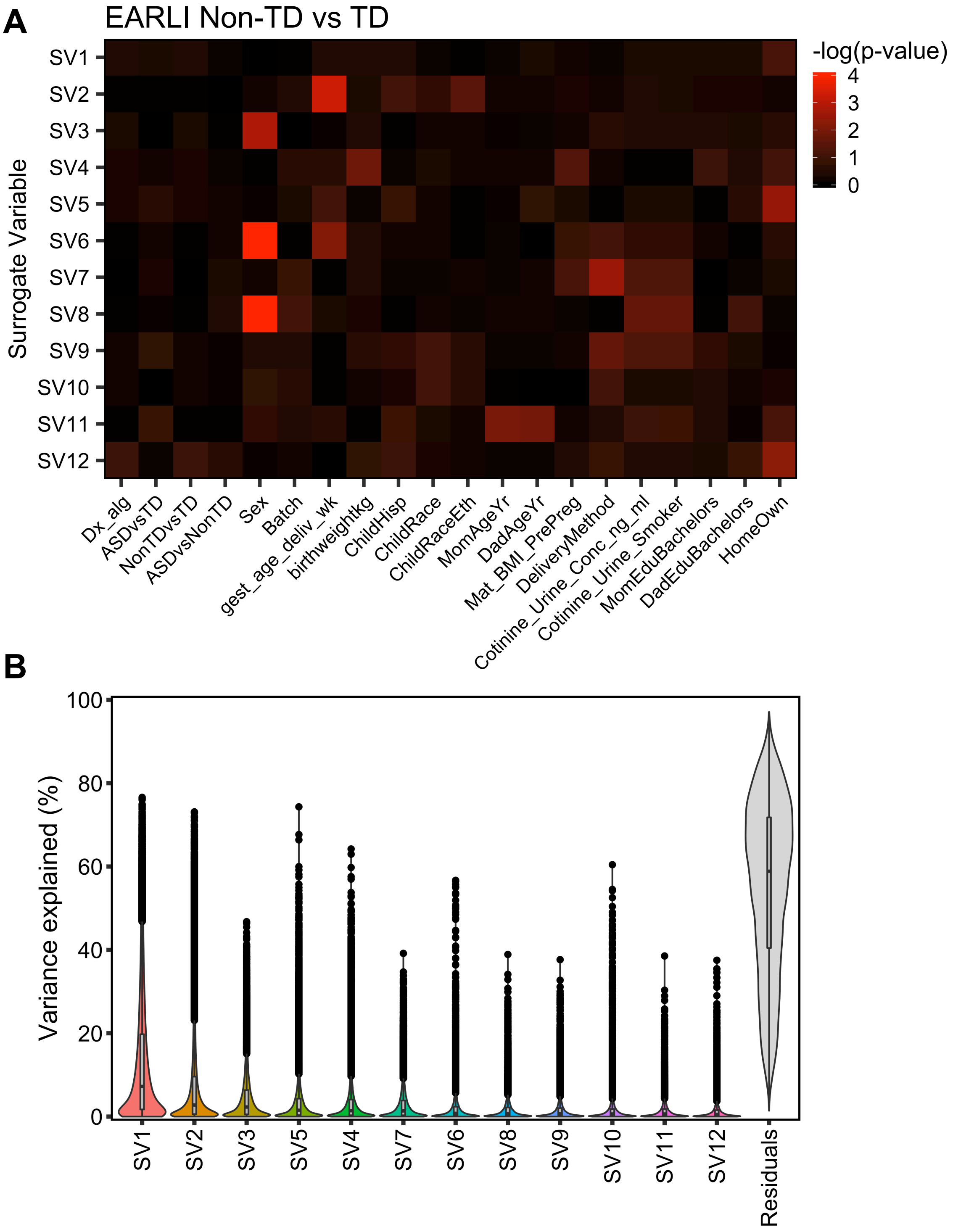


**Supplemental Figure 4. Surrogate variable analysis in EARLI subjects for the Non-TD versus TD comparison.** (A) Association of surrogate variables with covariates using linear regression. (B) Proportion of variance in expression of each probe explained by each surrogate variable, sorted by median variance explained.

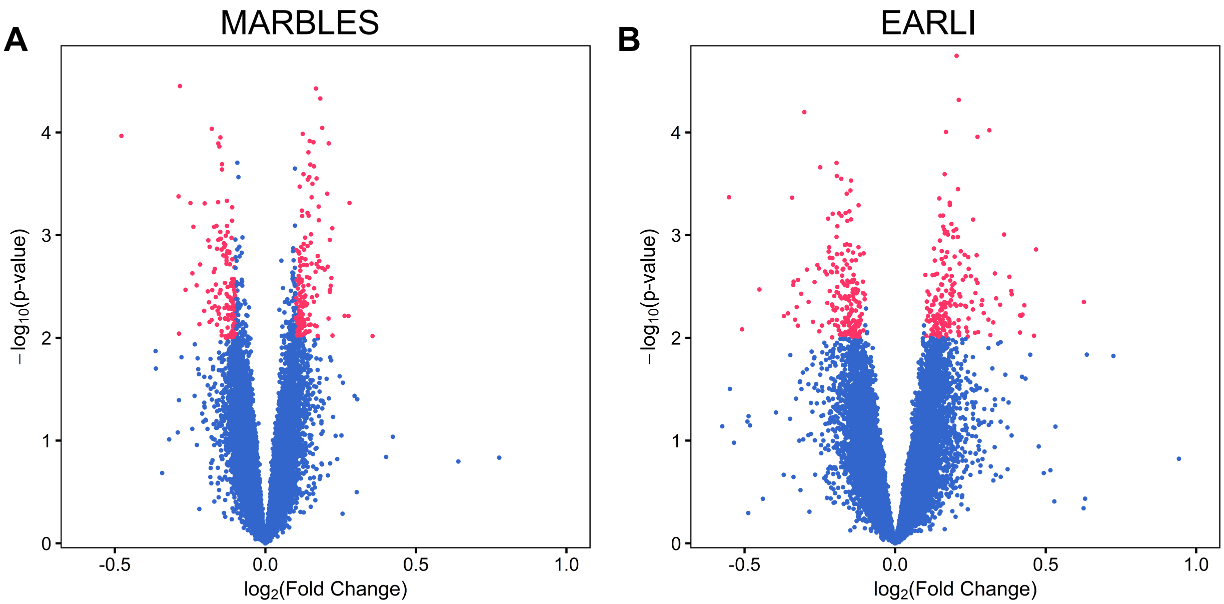


**Supplemental Figure 5. Identification of ASD-associated differentially-expressed genes in cord blood within each study.** Gene expression in umbilical cord blood samples from subjects with typical development or those diagnosed with ASD at age 3 was assessed by expression microarray. SVA was performed to control for technical and biological variables including sex and array batch. (A) Identification of 291 differentially-expressed genes in the MARBLES study (295 probes; $\log_2$(fold change) > 0.1, $p < 0.01$; TD $n = 77$, 40 male/37 female; ASD $n = 41$, 30 male/11 female). (B) Identification of 386 differentially-expressed genes in the EARLI study (392 probes; $\log_2$(fold change) > 0.1, $p < 0.01$; TD $n = 43$, 19 male/24 female; ASD $n = 18$, 13 male/5 female).

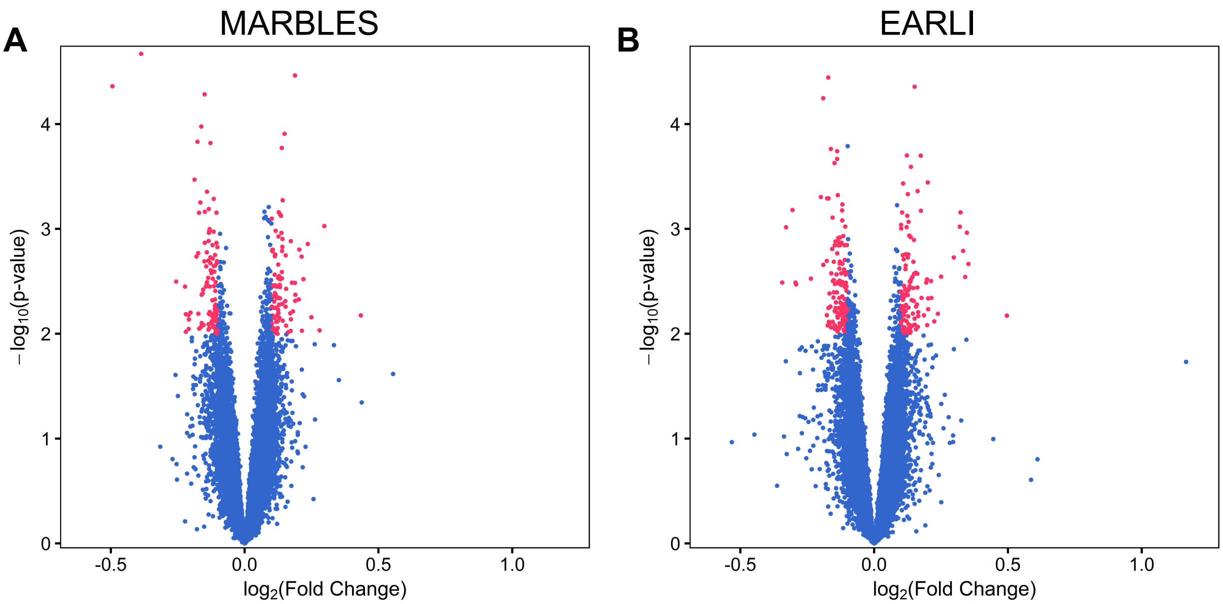


**Supplemental Figure 6. Identification of Non-TD-associated differentially-expressed genes in cord blood within each study.** Gene expression in umbilical cord blood samples from subjects with typical development or those diagnosed with Non-TD at age 3 was assessed by expression microarray. SVA was performed to control for technical and biological variables including sex and array batch. (A) Identification of 201 differentially-expressed genes in the MARBLES study (208 probes; $\log_2$(fold change) > 0.1, $p < 0.01$; TD $n = 77$, 40 male/37 female; Non-TD $n = 44$, 27 male/17 female). (B) Identification of 386 differentially-expressed genes in the EARLI study (392 probes; $\log_2$(fold change) > 0.1, $p < 0.01$; TD $n = 43$, 19 male/24 female; Non-TD $n = 48$, 23 male/25 female).

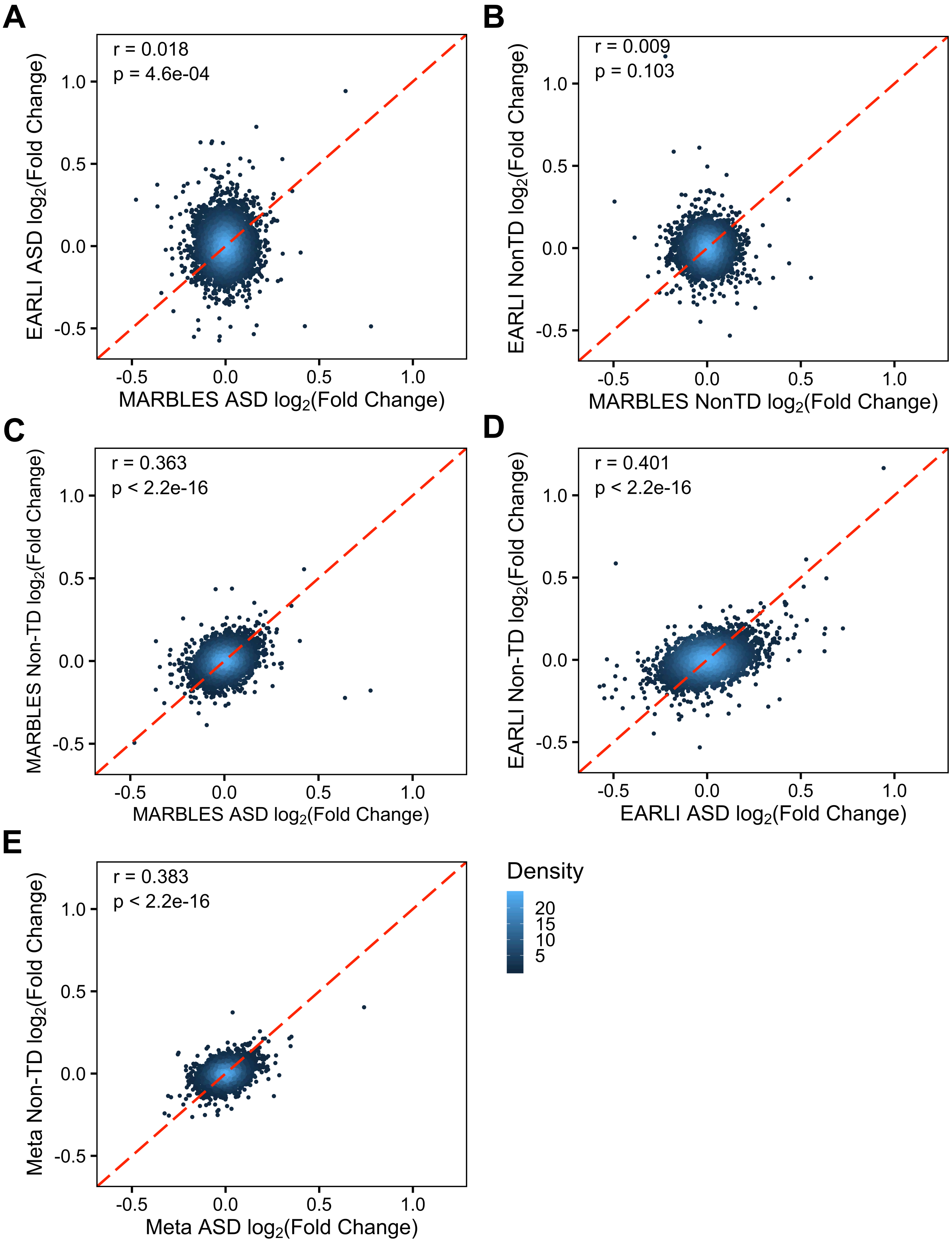


**Supplemental Figure 7. Correlations between ASD and Non-TD expression differences in MARBLES and EARLI subjects.** $log_2$(Fold Change) relative to TD expression is plotted for each probe, along with Pearson's correlation coefficients and p-values, for (A) ASD in MARBLES vs EARLI, (B) Non-TD in MARBLES vs EARLI, (C) ASD vs Non-TD in MARBLES, (D) ASD vs Non-TD in EARLI, or (E) ASD vs Non-TD in meta-analysis. Points are colored by density.

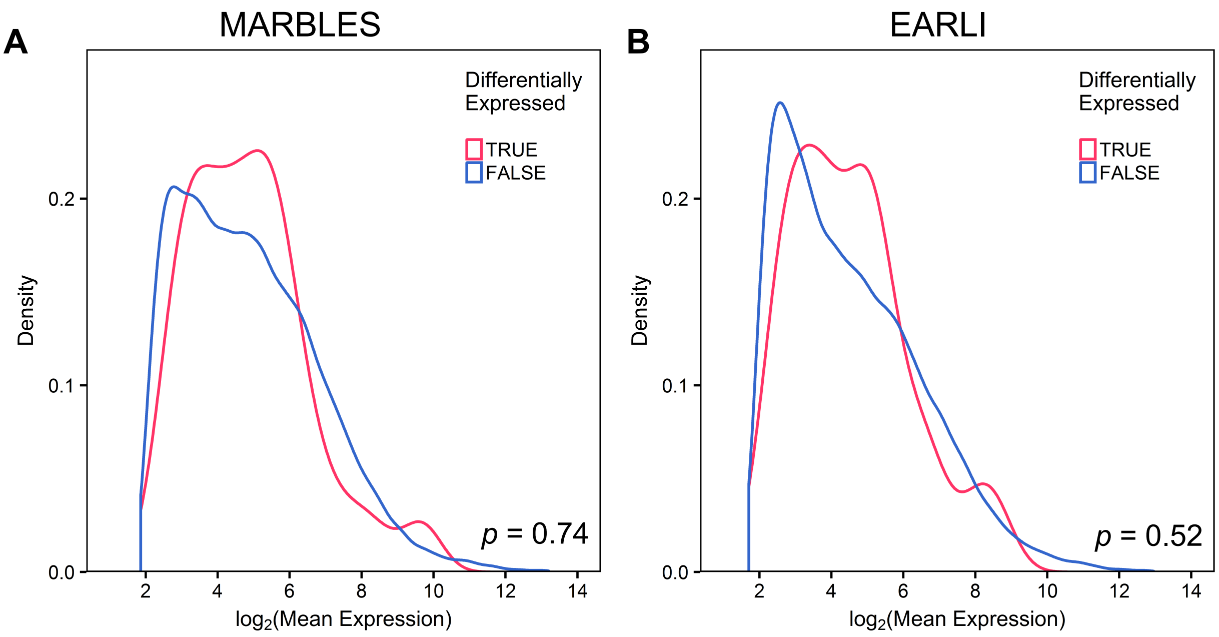


**Supplemental Figure 8. Expression level distribution of meta-analysis ASD versus TD differential probes is similar to non-differential probes.** Mean expression of meta-analysis ASD-associated probes was compared to probes with no difference in samples from the (A) MARBLES and (B) EARLI studies. Significance was assessed by comparing the median expression of differentially-expressed probes to the distribution of median expression of 10,000 equal-sized sets of randomly-sampled probes (MARBLES: differential = 4.70, non-differential = 4.64, $p$ = 0.74; EARLI: differential = 4.34, non-differential = 4.19, $p$ = 0.52).

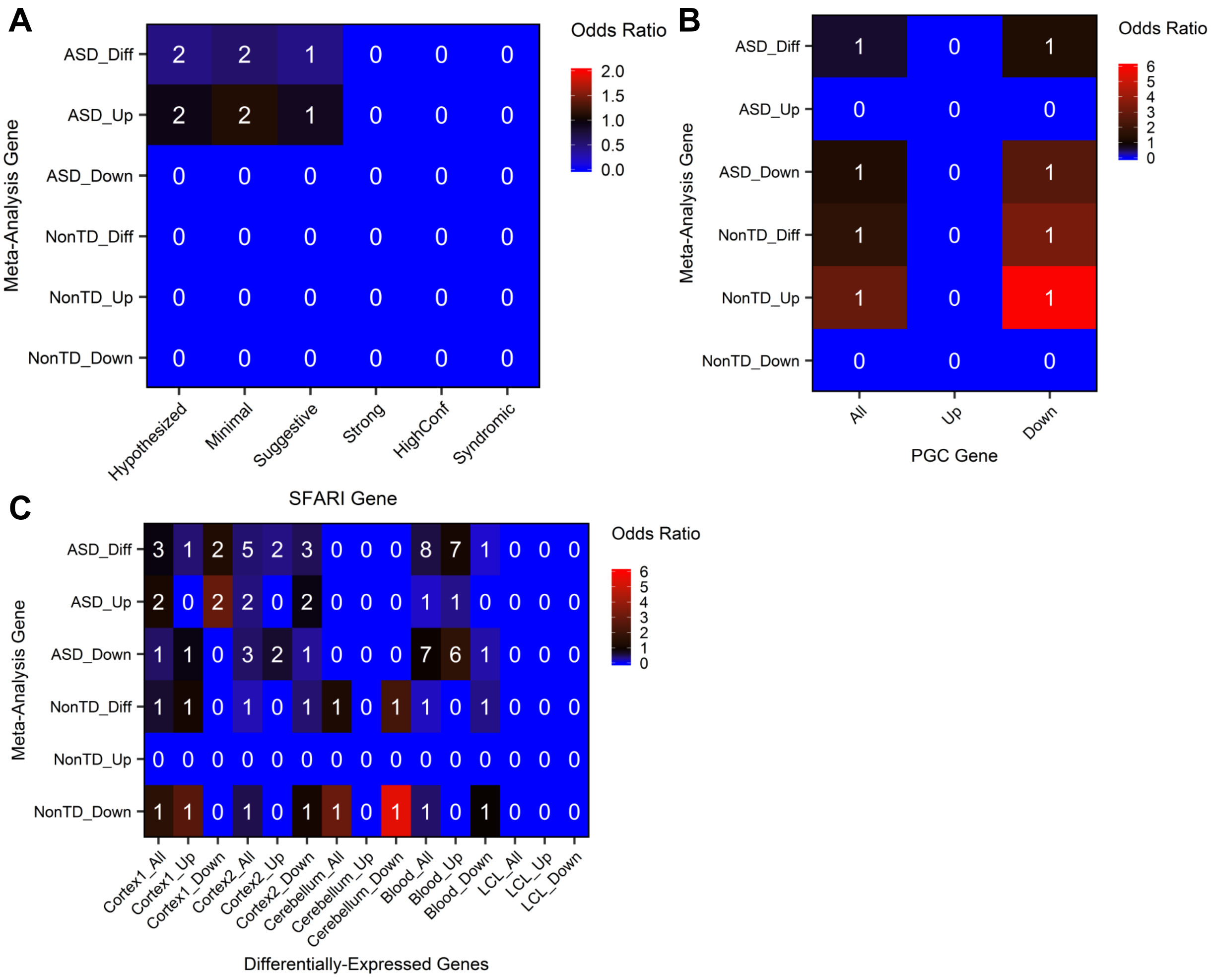


**Supplemental Figure 9. Cord blood differentially-expressed genes are not enriched for ASD-associated gene sets.** Differentially-expressed genes were overlapped with (A) putative ASD risk genes from SFARI gene by evidence level, (B) genes near ASD-associated SNPs from the psychiatric genomics consortium (PGC) by risk direction, or (C) differentially-expressed genes identified in ASD patients by tissue type. Heatmaps show number of overlapping genes and are colored by enrichment odds ratio. Significance was determined with Fisher's exact test (* FDR *q*-value < 0.05).

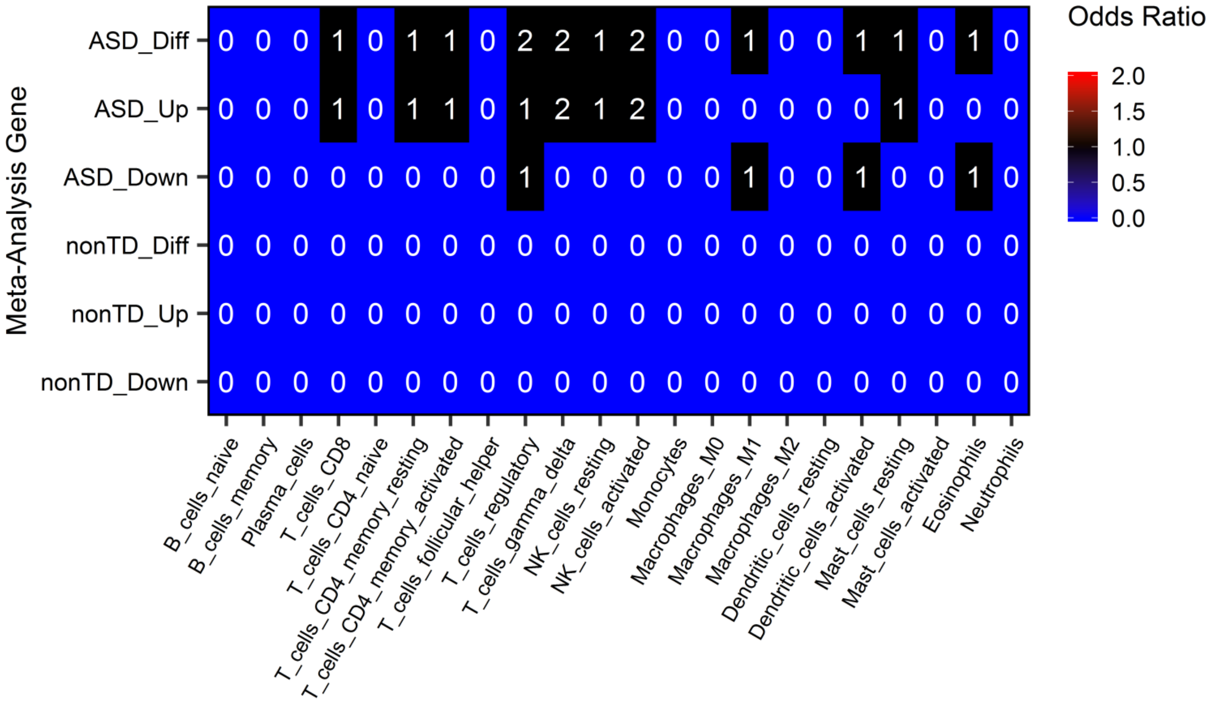


**Supplemental Figure 10. Cord blood differentially-expressed genes are depleted for blood cell-specific genes.** Differentially-expressed genes were overlapped with genes upregulated in purified cell types from peripheral blood (Newman et al. 2015). Heatmaps show number of overlapping genes and are colored by enrichment odds ratio. Significance was determined with Fisher's exact test (* FDR *q*-value < 0.05).

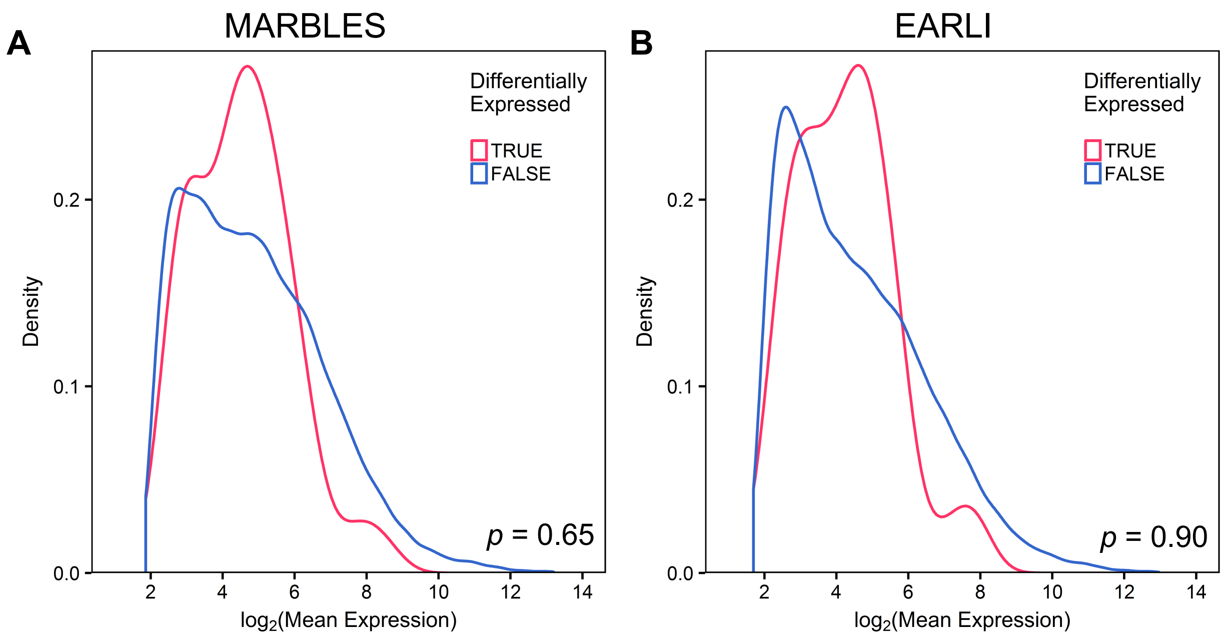


**Supplemental Figure 11. Expression level distribution of meta-analysis Non-TD versus TD differential probes is similar to non-differential probes.** Mean expression of meta-analysis Non-TD-associated probes was compared to probes with no difference in samples from the (A) MARBLES and (B) EARLI studies. Significance was assessed by comparing the median expression of differentially-expressed probes to the distribution of median expression of 10,000 equal-sized sets of randomly-sampled probes (MARBLES: differential = 4.48, non-differential = 4.64, $p$ = 0.65; EARLI: differential = 4.15, non-differential = 4.20, $p$ = 0.90).

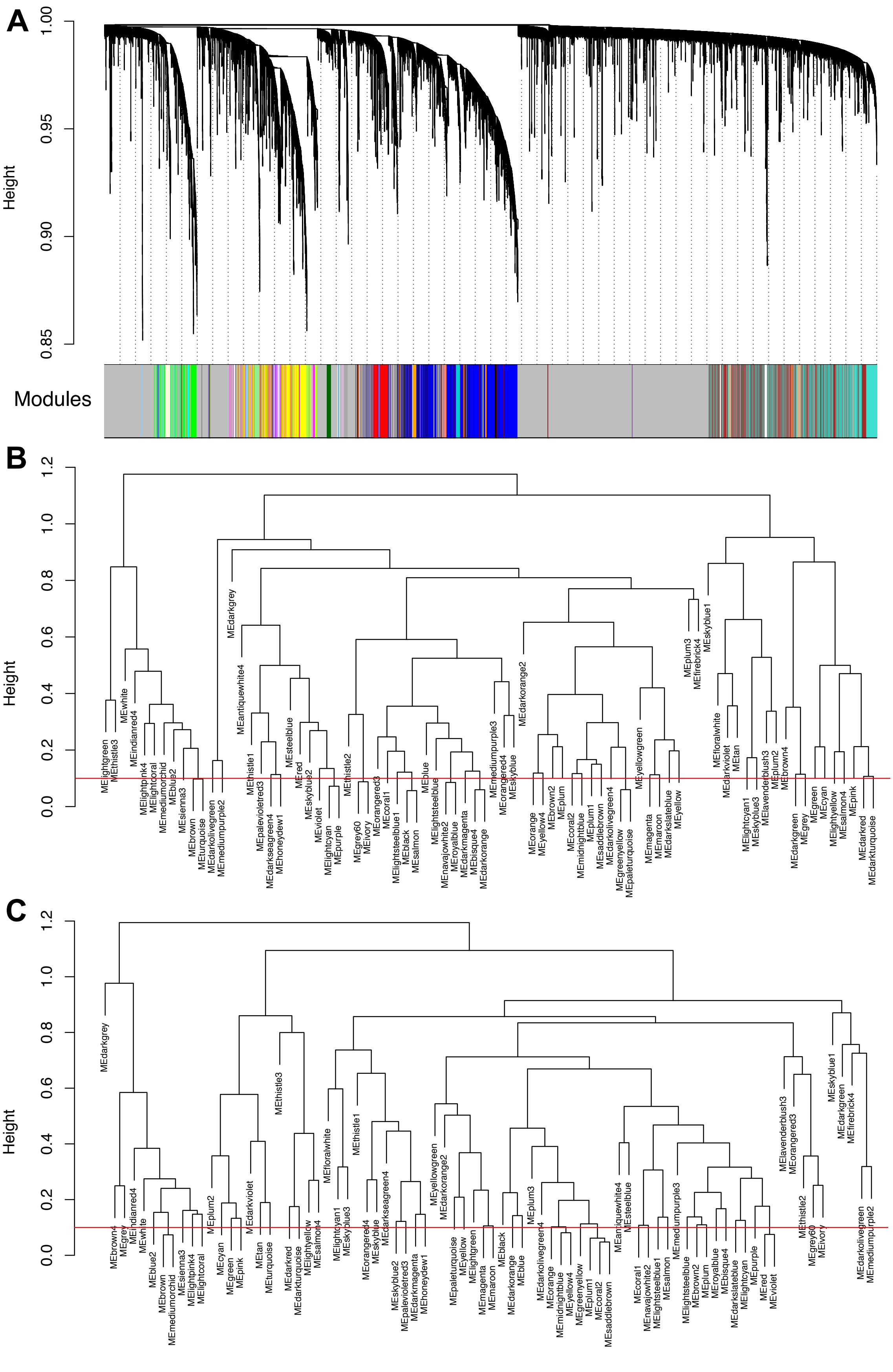
A
Height
1.00
0.95
0.90
0.85
Modules
B
Height
1.2
1.0
0.8
0.6
0.4
0.2
0.0
MElightgreen
MEthistle3
MEwhite
MEindianred4
MElightpink4
MElightcoral
MEmediumorchid
MEblue2
MEsienna3
MEbrown
MEturquoise
MEdarkolivegreen
MEmediumpurple2
MEdarkgrey
MEantiquewhite4
MEthistle1
MEpalevioletred3
MEdarkseagreen4
MEhoneydew1
MEsteelblue
MEred
MEskyblue2
MEviolet
MElightcyan
MEpurple
MEthistle2
MEgrey60
MEivory
MEorangered3
MEcoral1
MElightsteelblue1
MEblack
MEsalmon
MEblue
MElightsteelblue
MEnavajowhite2
MEroyalblue
MEdarkmagenta
MEbisque4
MEdarkorange
MEmediumpurple3
MEorangered4
MEskyblue
MEdarkorange2
MEorange
MEyellow4
MEbrown2
MEplum
MEcoral2
MEmidnightblue
MEplum1
MEsaddlebrown
MEdarkolivegreen4
MEgreenyellow
MEpaleturquoise
MEyellowgreen
MEmagenta
MEmaroon
MEdarkslateblue
MEyellow
MEplum3
MEfirebrick4
MEskyblue1
MEfloralwhite
MEdarkviolet
MEtan
MElightcyan1
MEskyblue3
MElavenderblush3
MEplum2
MEbrown4
MEdarkgreen
MEgrey
MEgreen
MEcyan
MElightyellow
MEsalmon4
MEpink
MEdarkred
MEdarkturquoise
C
Height
1.2
1.0
0.8
0.6
0.4
0.2
0.0
MEdarkgrey
MEbrown4
MEgrey
MEindianred4
MEwhite
MEblue2
MEbrown
MEmediumorchid
MEsienna3
MElightpink4
MElightcoral
MEplum2
MEcyan
MEgreen
MEpink
MEdarkviolet
MEtan
MEturquoise
MEthistle3
MEdarkred
MEdarkturquoise
MElightyellow
MEsalmon4
MEfloralwhite
MElightcyan1
MEskyblue3
MEthistle1
MEorangered4
MEskyblue
MEdarkseagreen4
MEskyblue2
MEpalevioletred3
MEdarkmagenta
MEhoneydew1
MEyellowgreen
MEdarkorange2
MEpaleturquoise
MEyellow
MElightgreen
MEmagenta
MEmaroon
MEdarkorange
MEblack
MEblue
MEplum3
MEdarkolivegreen4
MEorange
MEmidnightblue
MEyellow4
MEgreenyellow
MEplum1
MEcoral2
MEsaddlebrown
MEantiquewhite4
MEsteelblue
MEcoral1
MEnavajowhite2
MElightsteelblue1
MEsalmon
MEmediumpurple3
MElightsteelblue
MEbrown2
MEplum
MEroyalblue
MEbisque4
MEdarkslateblue
MElightcyan
MEpurple
MEred
MEviolet
MElavenderblush3
MEorangered3
MEthistle2
MEgrey60
MEivory
MEskyblue1
MEdarkgreen
MEfirebrick4
MEdarkolivegreen
MEmediumpurple2

**Supplemental Figure 12. Consensus coexpression modules identified in MARBLES and EARLI.** (A) Gene dendrogram clustered by normalized intensity with identified consensus modules. (B) Module dendrogram clustered by consensus module eigengenes in MARBLES subjects. (C) Module dendrogram clustered by consensus module eigengenes in EARLI subjects.

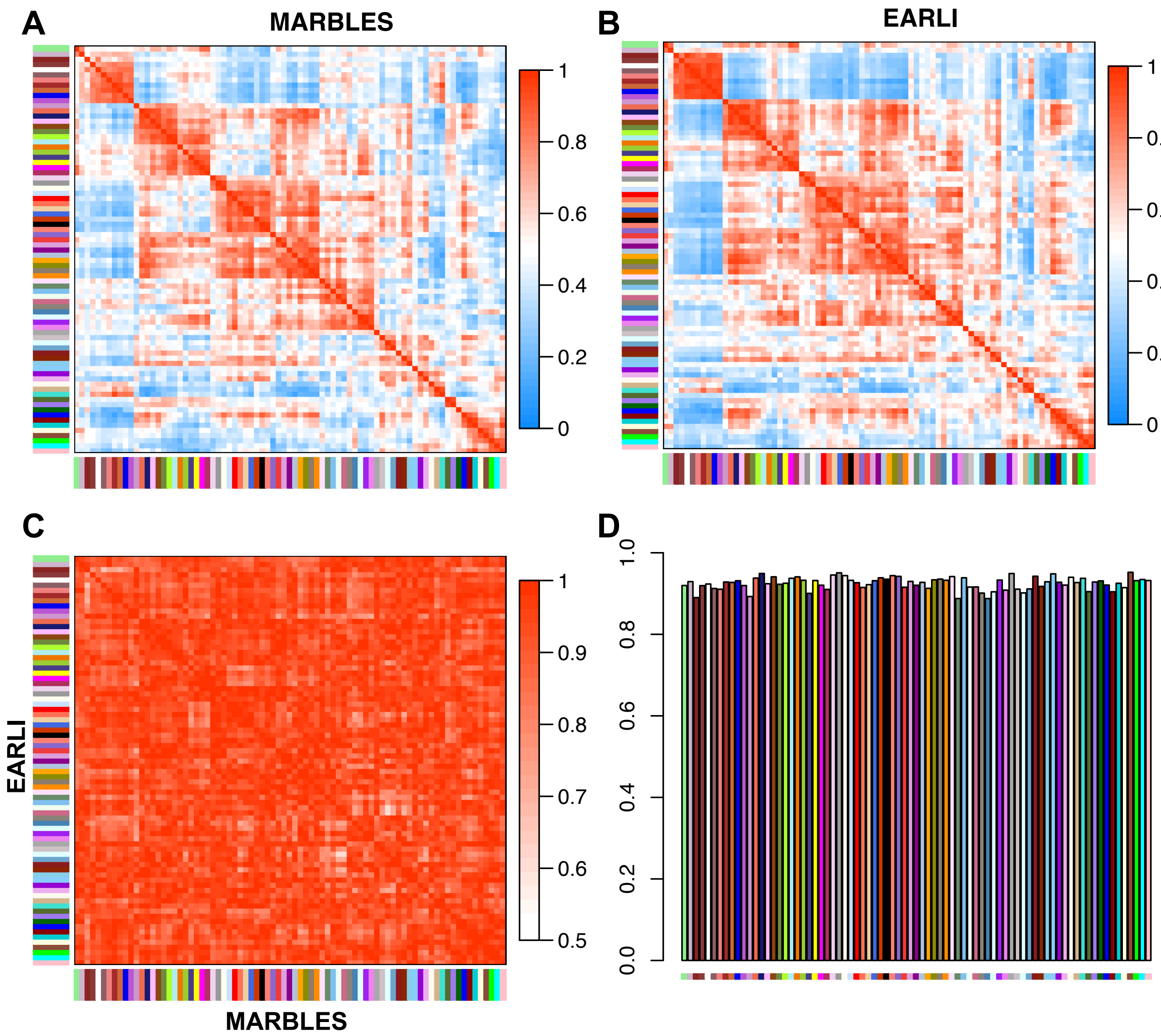


**Supplemental Figure 13. Consensus module eigengene networks are preserved between MARBLES and EARLI subjects.** (A) Correlation between consensus module eigengenes in

MARBLES subjects. (B) Correlation between consensus module eigengenes in EARLI subjects. (C) Pairwise preservation of consensus module eigengene correlations between MARBLES and EARLI subjects. (D) Mean preservation of consensus module eigengene correlations between MARBLES and EARLI subjects. Overall preservation = 0.93.

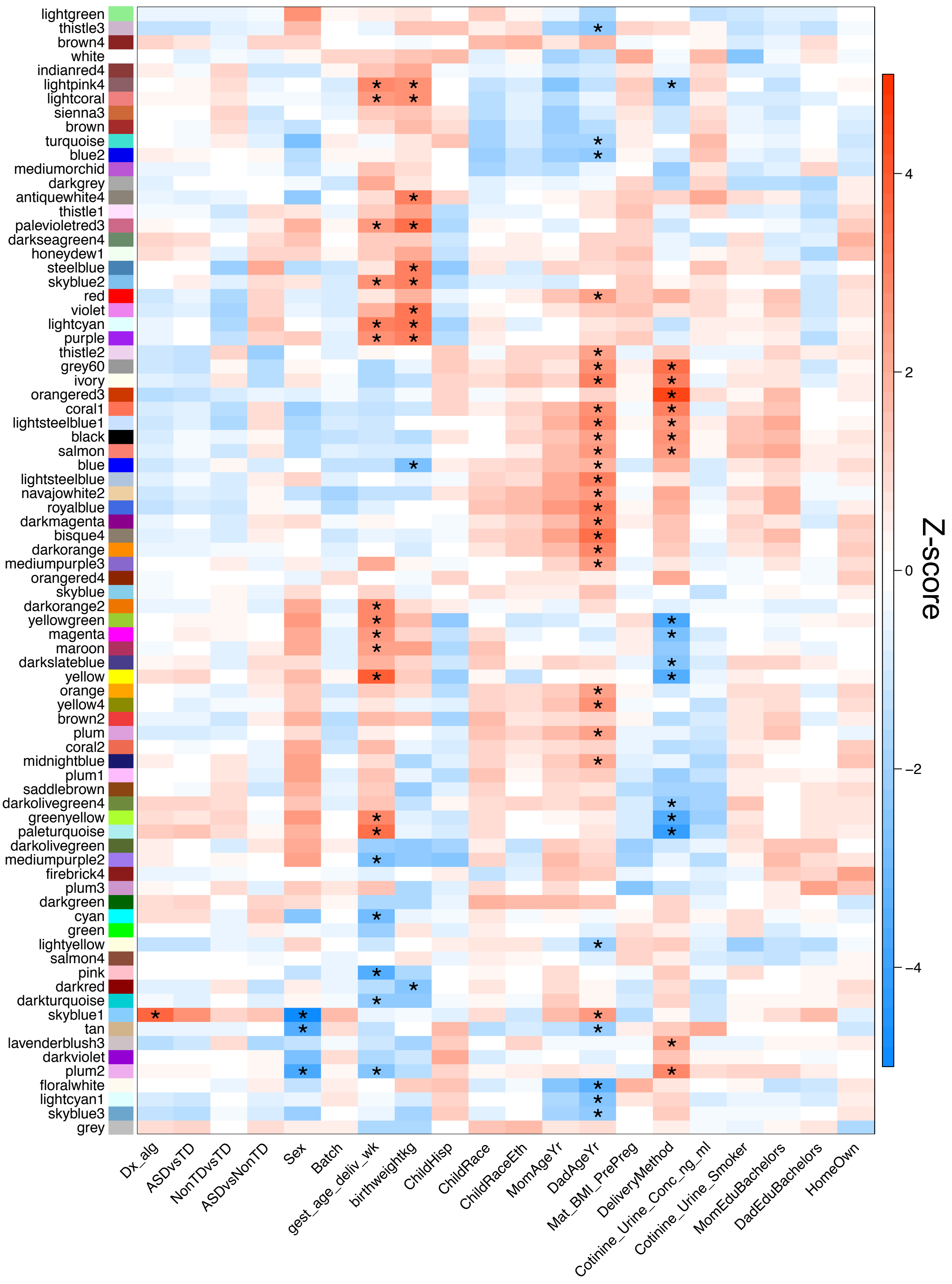
lightgreen
thistle3
brown4
white
indianred4
lightpink4
lightcoral
sienna3
brown
turquoise
blue2
mediumorchid
darkgrey
antiquewhite4
thistle1
palevioletred3
darkseagreen4
honeydew1
steelblue
skyblue2
red
violet
lightcyan
purple
thistle2
grey60
ivory
orangered3
coral1
lightsteelblue1
black
salmon
blue
lightsteelblue
navajowhite2
royalblue
darkmagenta
bisque4
darkorange
mediumpurple3
orangered4
skyblue
darkorange2
yellowgreen
magenta
maroon
darkslateblue
yellow
orange
yellow4
brown2
plum
coral2
midnightblue
plum1
saddlebrown
darkolivegreen4
greenyellow
paleturquoise
darkolivegreen
mediumpurple2
firebrick4
plum3
darkgreen
cyan
green
lightyellow
salmon4
pink
darkred
darkturquoise
skyblue1
tan
lavenderblush3
darkviolet
plum2
floralwhite
lightcyan1
skyblue3
grey
Dx_alg
ASDvsTD
NonTDvsTD
ASDvsNonTD
Sex
Batch
gest_age_deliv_wk
birthweightkg
ChildHisp
ChildRace
ChildRaceEth
MomAgeYr
DadAgeYr
Mat_BMI_PrePreg
DeliveryMethod
Cotinine_Urine_Conc_ng_ml
Cotinine_Urine_Smoker
MomEduBachelors
DadEduBachelors
HomeOwn
Z-score
4
2
0
−2
−4

1

**Supplemental Figure 14. Consensus modules are correlated with diagnosis and demographic factors in MARBLES subjects.** Heatmap of biweight midcorrelation Z-scores of module eigengenes with sample covariates (ASD $n$ = 41, Non-TD $n$ = 44, TD $n$ = 76). P-values were adjusted for all comparisons using the FDR method (* $q$ < 0.05).

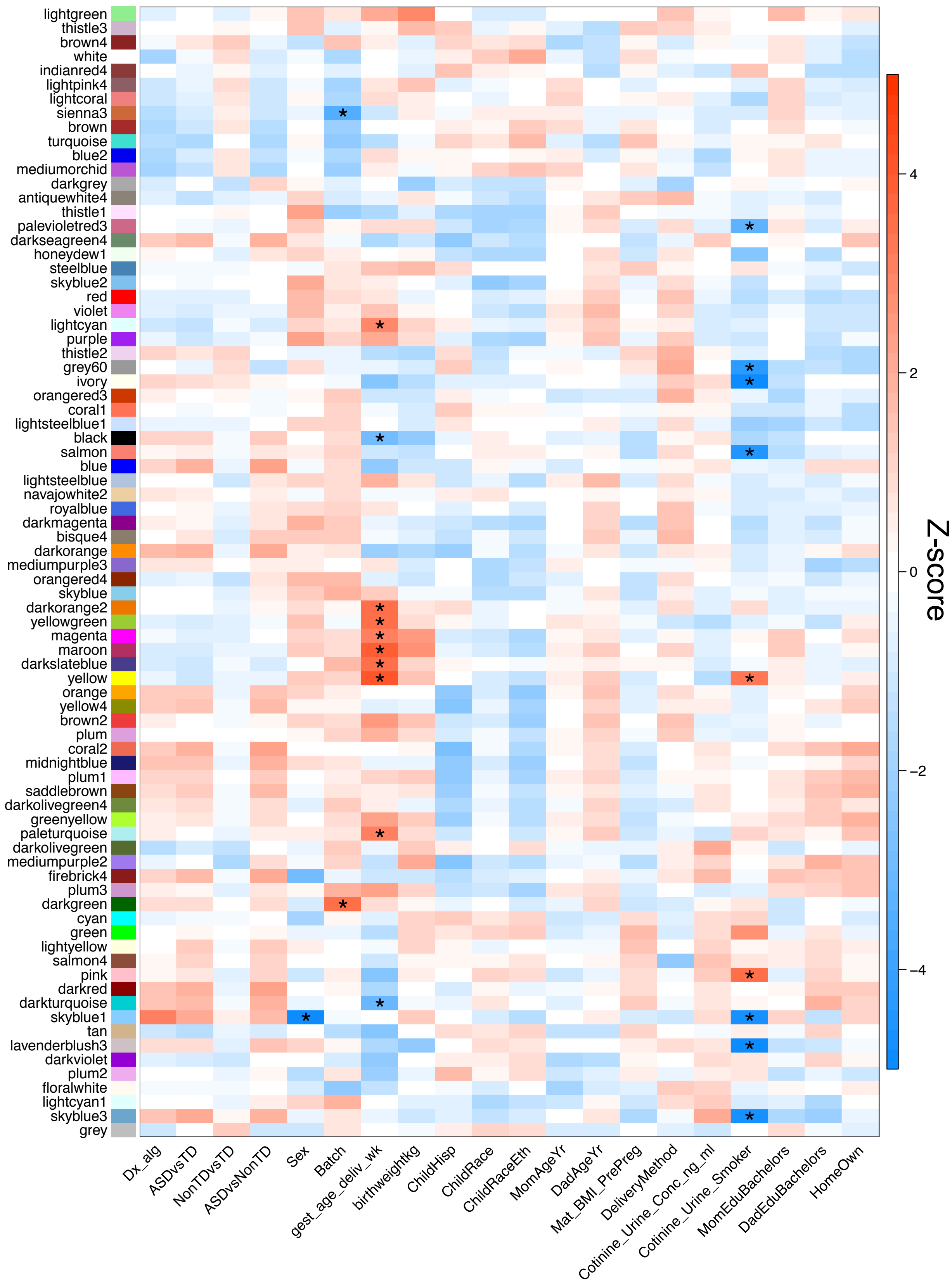
lightgreen
thistle3
brown4
white
indianred4
lightpink4
lightcoral
sienna3
brown
turquoise
blue2
mediumorchid
darkgrey
antiquewhite4
thistle1
palevioletred3
darkseagreen4
honeydew1
steelblue
skyblue2
red
violet
lightcyan
purple
thistle2
grey60
ivory
orangered3
coral1
lightsteelblue1
black
salmon
blue
lightsteelblue
navajowhite2
royalblue
darkmagenta
bisque4
darkorange
mediumpurple3
orangered4
skyblue
darkorange2
yellowgreen
magenta
maroon
darkslateblue
yellow
orange
yellow4
brown2
plum
coral2
midnightblue
plum1
saddlebrown
darkolivegreen4
greenyellow
paleturquoise
darkolivegreen
mediumpurple2
firebrick4
plum3
darkgreen
cyan
green
lightyellow
salmon4
pink
darkred
darkturquoise
skyblue1
tan
lavenderblush3
darkviolet
plum2
floralwhite
lightcyan1
skyblue3
grey
Dx_alg
ASDvsTD
NonTDvsTD
ASDvsNonTD
Sex
Batch
gest_age_deliv_wk
birthweightkg
ChildHisp
ChildRace
ChildRaceEth
MomAgeYr
DadAgeYr
Mat_BMI_PrePreg
DeliveryMethod
Cotinine_Urine_Conc_ng_ml
Cotinine_Urine_Smoker
MomEduBachelors
DadEduBachelors
HomeOwn
Z-score
4
2
0
−2
−4

**Supplemental Figure 15. Consensus modules are correlated with demographic factors in EARLI subjects.** Heatmap of biweight midcorrelation Z-scores of module eigengenes with sample covariates (ASD *n* = 19, Non-TD *n* = 47, TD *n* = 43). P-values were adjusted for all comparisons using the FDR method (* $q < 0.05$).

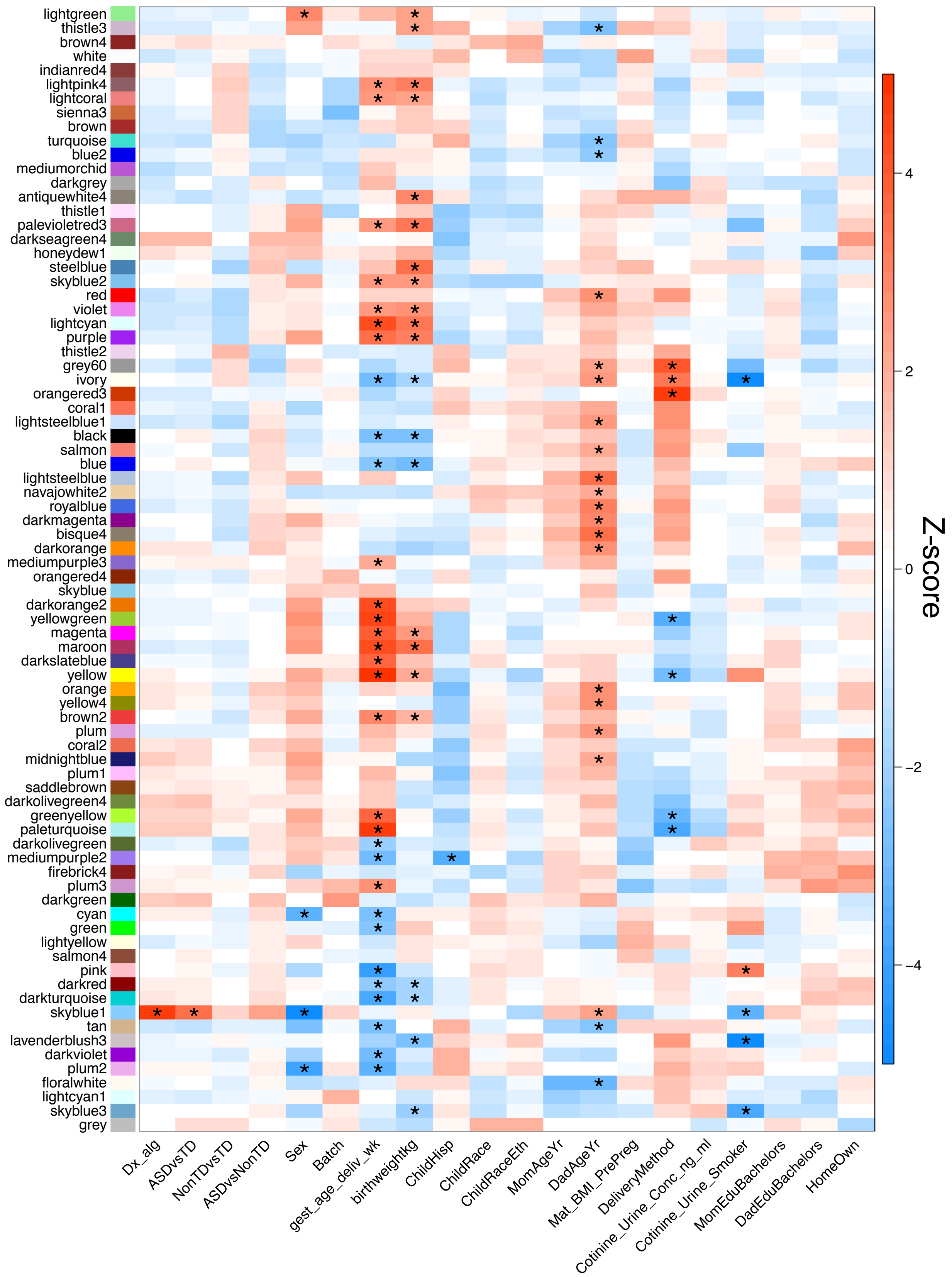
lightgreen
thistle3
brown4
white
indianred4
lightpink4
lightcoral
sienna3
brown
turquoise
blue2
mediumorchid
darkgrey
antiquewhite4
thistle1
palevioletred3
darkseagreen4
honeydew1
steelblue
skyblue2
red
violet
lightcyan
purple
thistle2
grey60
ivory
orangered3
coral1
lightsteelblue1
black
salmon
blue
lightsteelblue
navajowhite2
royalblue
darkmagenta
bisque4
darkorange
mediumpurple3
orangered4
skyblue
darkorange2
yellowgreen
magenta
maroon
darkslateblue
yellow
orange
yellow4
brown2
plum
coral2
midnightblue
plum1
saddlebrown
darkolivegreen4
greenyellow
paleturquoise
darkolivegreen
mediumpurple2
firebrick4
plum3
darkgreen
cyan
green
lightyellow
salmon4
pink
darkred
darkturquoise
skyblue1
tan
lavenderblush3
darkviolet
plum2
floralwhite
lightcyan1
skyblue3
grey
Dx_alg
ASDvsTD
NonTDvsTD
ASDvsNonTD
Sex
Batch
gest_age_deliv_wk
birthweightkg
ChildHisp
ChildRace
ChildRaceEth
MomAgeYr
DadAgeYr
Mat_BMI_PrePreg
DeliveryMethod
Cotinine_Urine_Conc_ng_ml
Cotinine_Urine_Smoker
MomEduBachelors
DadEduBachelors
HomeOwn
Z-score
4
2
0
−2
−4

**Supplemental Figure 16. Consensus modules are correlated with diagnosis and demographic factors in meta-analysis.** Heatmap of meta-analysis biweight midcorrelation Z-scores of module eigengenes with sample covariates (MARBLES: ASD *n* = 41, Non-TD *n* = 44, TD *n* = 76; EARLI: ASD *n* = 19, Non-TD *n* = 47, TD *n* = 43). Z-scores from the individual studies were combined using Stouffer's method with weights given by the square root of sample *n.* P-values were adjusted for all comparisons using the FDR method (* $q <$ 0.05).

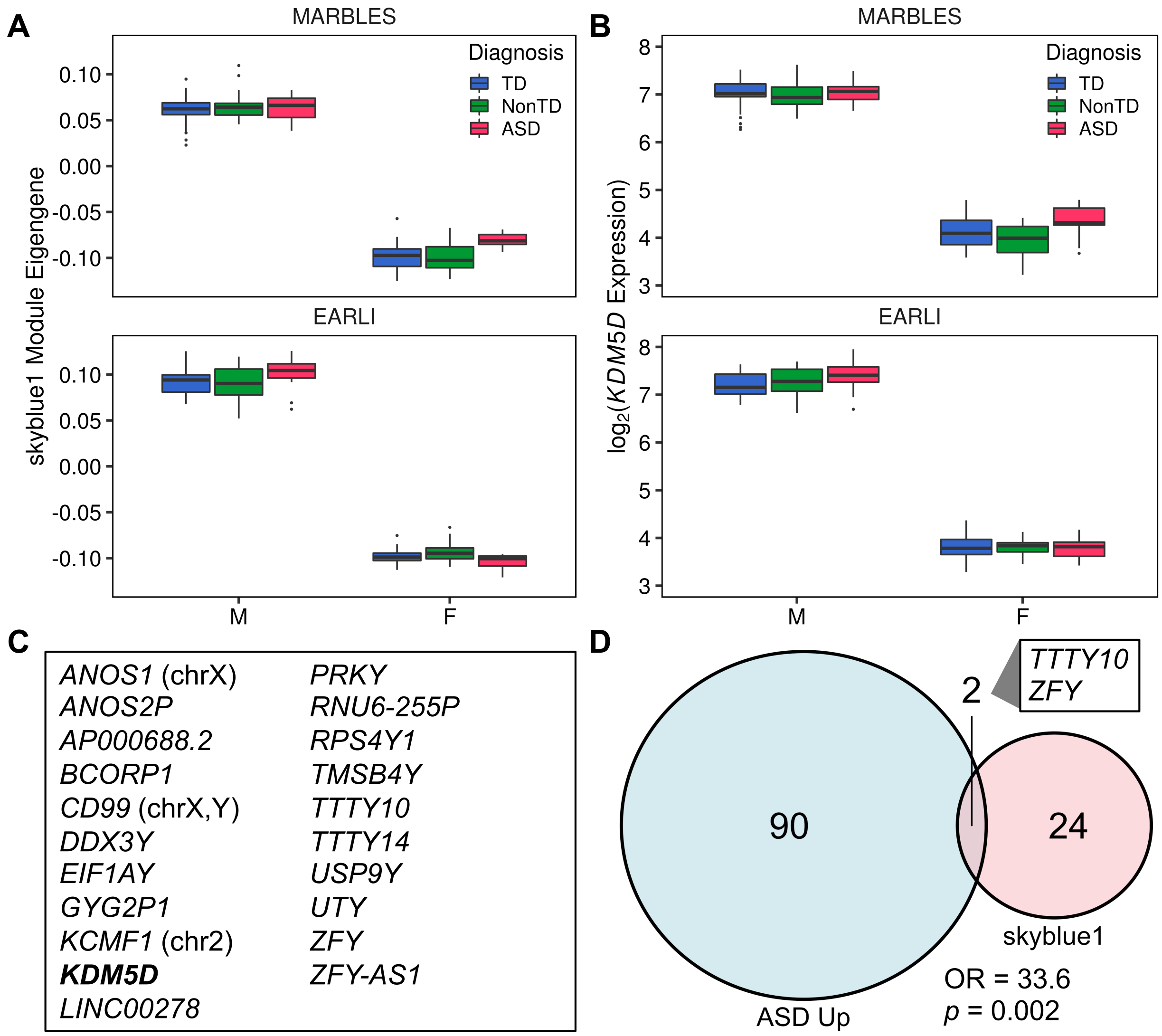

**Supplemental Figure 17. Skyblue1 module is specifically expressed in males and is enriched for genes upregulated in ASD.** Boxplots of (A) skyblue1 module eigengene or (B) Normalized $log_2$(expression) of skyblue1 hub gene *KDM5D* by diagnosis and sex. P-values for the skyblue1 module eigengene were adjusted for the total number of modules using the FDR method. (Meta-analysis ME ~ Diagnosis $q$ = 1.3E-4, ME ~ Sex $q$ = 3.4E-172; Meta-analysis *KDM5D* ~ Diagnosis $p$ = 1.2E-6, *KDM5D* ~ Sex $p$ = 6.4E-164). (C) Genes annotated to probes in the skyblue1 module. Genes are located on the Y chromosome unless otherwise indicated. (D) Overlap of skyblue1 module probes with probes upregulated in ASD meta-analysis. Overlap significance was assessed using Fisher's exact test.

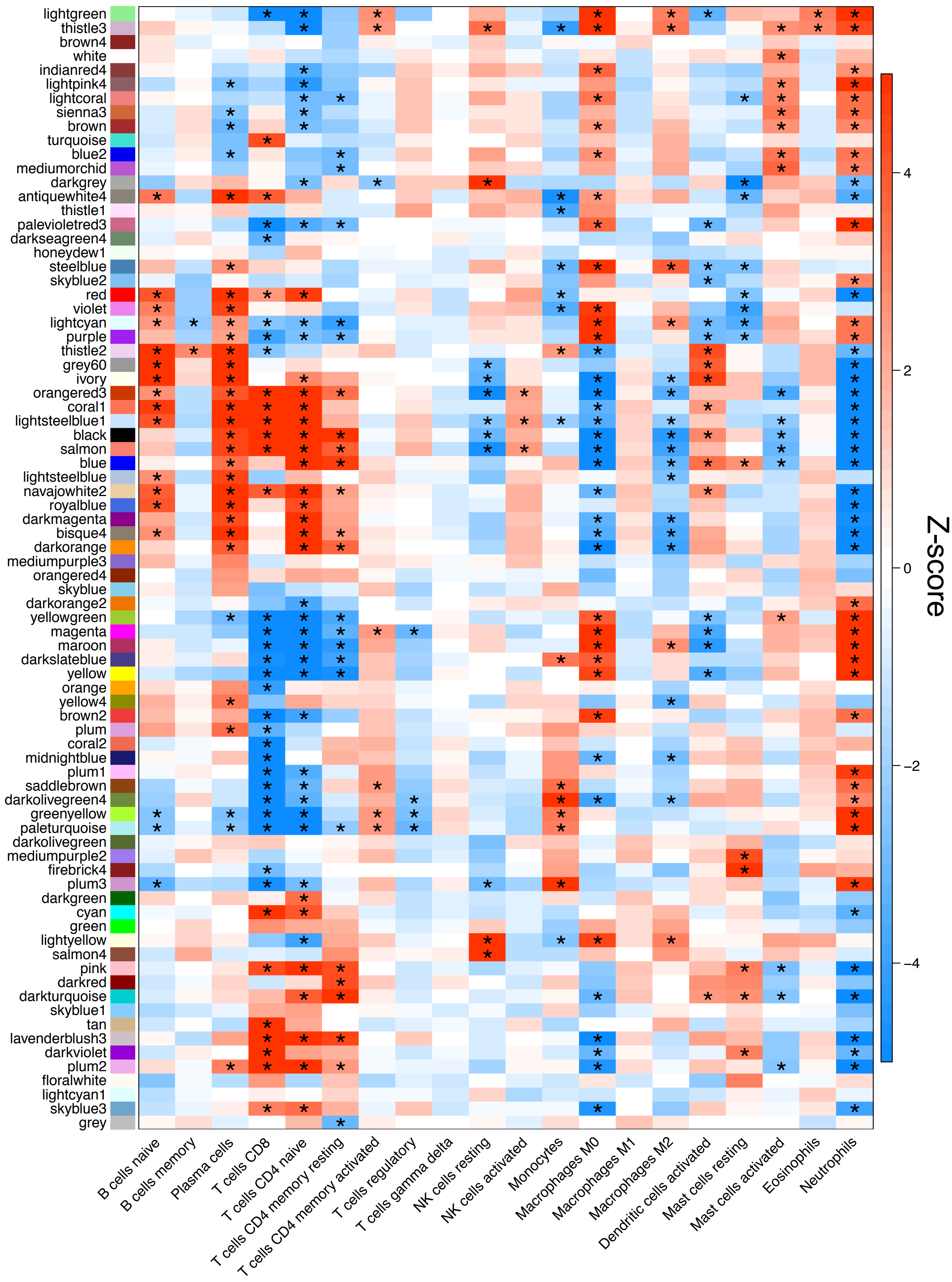
lightgreen
thistle3
brown4
white
indianred4
lightpink4
lightcoral
sienna3
brown
turquoise
blue2
mediumorchid
darkgrey
antiquewhite4
thistle1
palevioletred3
darkseagreen4
honeydew1
steelblue
skyblue2
red
violet
lightcyan
purple
thistle2
grey60
ivory
orangered3
coral1
lightsteelblue1
black
salmon
blue
lightsteelblue
navajowhite2
royalblue
darkmagenta
bisque4
darkorange
mediumpurple3
orangered4
skyblue
darkorange2
yellowgreen
magenta
maroon
darkslateblue
yellow
orange
yellow4
brown2
plum
coral2
midnightblue
plum1
saddlebrown
darkolivegreen4
greenyellow
paleturquoise
darkolivegreen
mediumpurple2
firebrick4
plum3
darkgreen
cyan
green
lightyellow
salmon4
pink
darkred
darkturquoise
skyblue1
tan
lavenderblush3
darkviolet
plum2
floralwhite
lightcyan1
skyblue3
grey
B cells naive
B cells memory
Plasma cells
T cells CD8
T cells CD4 naive
T cells CD4 memory resting
T cells CD4 memory activated
T cells regulatory
T cells gamma delta
NK cells resting
NK cells activated
Monocytes
Macrophages M0
Macrophages M1
Macrophages M2
Dendritic cells activated
Mast cells resting
Mast cells activated
Eosinophils
Neutrophils
Z-score
4
2
0
−2
−4

**Supplemental Figure 18. Consensus modules are strongly correlated with cell type proportions in meta-analysis.** Heatmap of meta-analysis biweight midcorrelation Z-scores of module eigengenes with cell type proporitions (MARBLES: ASD $n$ = 41, Non-TD $n$ = 44, TD $n$ = 76; EARLI: ASD $n$ = 19, Non-TD $n$ = 47, TD $n$ = 43). Z-scores from the individual studies were combined using Stouffer's method with weights given by the square root of sample $n$. P-values were adjusted for all comparisons using the FDR method (* $q < 0.05$).

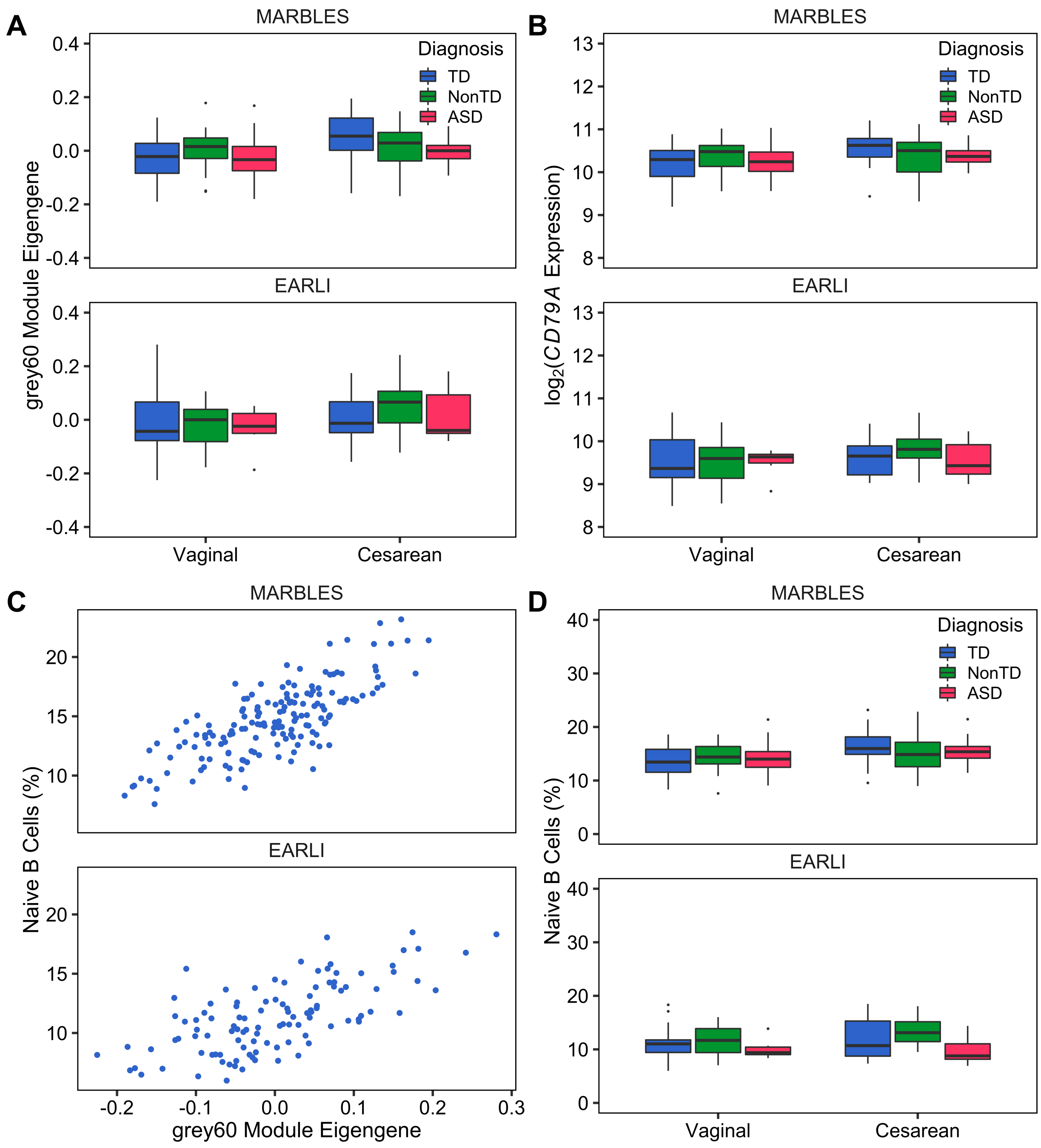


**Supplemental Figure 19. The B cell-associated grey60 module is upregulated during cesarean delivery and is downregulated in ASD.** Boxplots of (A) grey60 module eigengene or (B) Normalized $\log_2$(expression) of grey60 hub gene *CD79A* by diagnosis, delivery method, and study. (Meta-analysis ME ~ ASD vs Non-TD $p = 0.048$, ME ~ Delivery Method $p = 4.7E\text{-}5$; Meta-analysis *CD79A* ~ ASD vs Non-TD $p = 0.047$, *CD79A* ~ Delivery

Method *p* =4.2E-4). The grey60 module contains the genes *IGLV1-40* which is upregulated in both ASD and Non-TD compared to TD, and *IGLC2* which is upregulated in ASD vs TD. (C) Scatterplot of grey60 module eigengene compared to estimated proportion of naïve B cells (Meta-analysis B cells ~ ME *p* = 8.8E-46). (D) Boxplot of naïve B cells by diagnosis, delivery method, and study (Meta-analysis B cells ~ ASD vs Non-TD *p* = 0.020, B cells ~ Delivery Method *p* = 1.7E-4).
